## Supplementary figures and images for "Adenomatous Polyposis Coli Loss Controls Cell Cycle Regulators and Response to Paclitaxel"

### Supplemental Figure 1A

Supplementary Figure 1A. Cell cycle

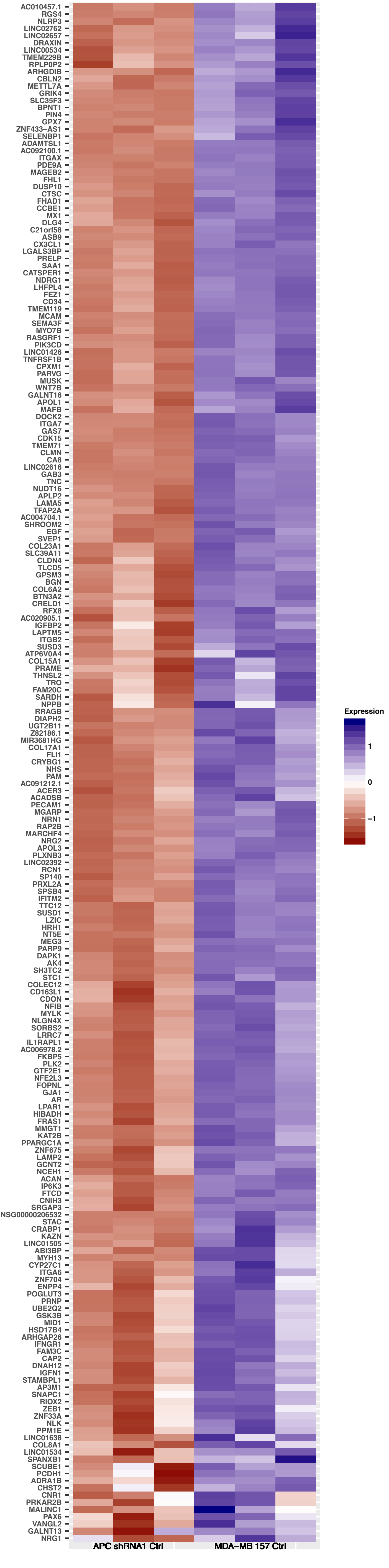

### Supplemental Figure 2A

Supplementary Figure 2A. Cell cycle

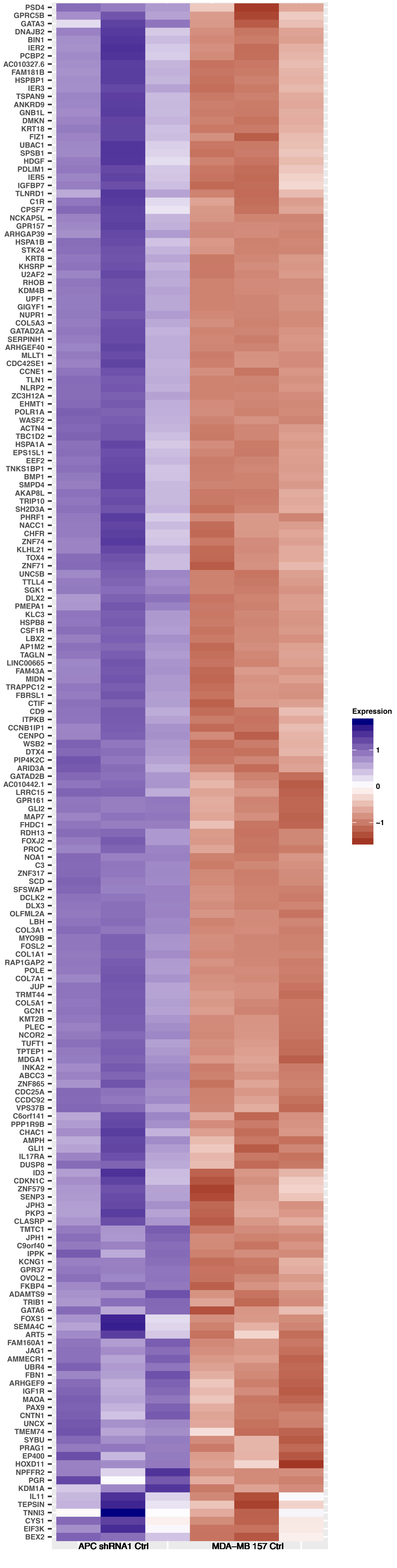
