## Supplemental Figure 1B for "Adenomatous Polyposis Coli Loss Controls Cell Cycle Regulators and Response to Paclitaxel"

Supplemental Figure 1B. Down-regulated in APC shRNA1 cells

| ensembl_gene_id | hgnc | entrezgene_id | clusterOrder | baseMean | log2FoldChange | lfcSE | stat | pvalue | description |
|---|---|---|---|---|---|---|---|---|---|
| **ENSG00000251257** | AC010457.1 (novel transcript) | NA | 214 | 46.40621 | -4.140618 | 0.5478044 | -7.55857 | 4.08E-14 | LncRNA |
| **ENSG00000117152** | RGS4 | 5999 | 213 | 2.886760776 | -5.961099825 | 1.354142335 | -4.402122044 | 1.07E-05 | regulator of G protein signaling 4 [Source:HGNC Symbol;Acc:HGNC:10000] |
| **ENSG00000162711** | NLRP3 | 114548 | 212 | 26.33632458 | -2.932799097 | 0.4072063 | -7.202243909 | 5.92E-13 | NLR family pyrin domain containing 3 [Source:HGNC Symbol;Acc:HGNC:16400] |
| **ENSG00000250303** | LINC02762 | NA | 211 | 45.7454375 | -1.587040232 | 0.266763927 | -5.949231033 | 2.69E-09 | long intergenic non-protein coding RNA 2762 [Source:HGNC Symbol;Acc:HGNC:27443] |
| **ENSG00000242147** | LINC02657 | NA | 210 | 83.16373373 | -1.391442155 | 0.258073612 | -5.391648317 | 6.98E-08 | long intergenic non-protein coding RNA 2657 [Source:HGNC Symbol;Acc:HGNC:54143] |
| **ENSG00000162490** | DRAXIN | 374946 | 209 | 29.47874841 | -2.497354223 | 0.352701197 | -7.080651397 | 1.43E-12 | dorsal inhibitory axon guidance protein [Source:HGNC Symbol;Acc:HGNC:25054] |
| **ENSG00000253394** | LINC00534 | NA | 208 | 16.25941805 | -0.925089188 | 0.315367313 | -2.933370548 | 0.003353034 | long intergenic non-protein coding RNA 534 [Source:HGNC Symbol;Acc:HGNC:43643] |
| **ENSG00000198133** | TMEM229B | 161145 | 207 | 35.28219165 | -1.325904817 | 0.297233501 | -4.460818894 | 8.16E-06 | transmembrane protein 229B [Source:HGNC Symbol;Acc:HGNC:20130] |
| **ENSG00000243742** | RPLP0P2 | NA | 206 | 26.71983697 | -2.15901419 | 0.371987804 | -5.803991877 | 6.48E-09 | ribosomal protein lateral stalk subunit P0 pseudogene 2 [Source:HGNC Symbol;Acc:HGNC:17960] |
| **ENSG00000111348** | ARHGDIB | 397 | 205 | 121.6175867 | -1.625309897 | 0.208135999 | -7.808884108 | 5.77E-15 | Rho GDP dissociation inhibitor beta [Source:HGNC Symbol;Acc:HGNC:679] |
| **ENSG00000141668** | CBLN2 | 147381 | 204 | 3.558657039 | -3.755870532 | 0.978170698 | -3.839688248 | 0.000123191 | cerebellin 2 precursor [Source:HGNC Symbol;Acc:HGNC:1544] |
| **ENSG00000185432** | METTL7A | 25840 | 203 | 133.7953679 | -1.936537482 | 0.192949807 | -10.03648312 | 1.05E-23 | methyltransferase like 7A [Source:HGNC Symbol;Acc:HGNC:24550] |
| **ENSG00000149403** | GRIK4 | 2900 | 202 | 65.89392186 | -1.283414898 | 0.177651248 | -7.224350584 | 5.04E-13 | glutamate ionotropic receptor kainate type subunit 4 [Source:HGNC Symbol;Acc:HGNC:4582] |
| **ENSG00000183780** | SLC35F3 | 148641 | 201 | 341.3943602 | -0.796980835 | 0.123802016 | -6.437543296 | 1.21E-10 | solute carrier family 35 member F3 [Source:HGNC Symbol;Acc:HGNC:23616] |
| **ENSG00000162813** | BPNT1 | 10380 | 200 | 187.8919417 | -1.539891345 | 0.181974594 | -8.462122717 | 2.63E-17 | 3'(2'), 5'-bisphosphate nucleotidase 1 [Source:HGNC Symbol;Acc:HGNC:1096] |
| **ENSG00000102309** | PIN4 | 5303 | 199 | 322.2198344 | -1.103333245 | 0.20165113 | -5.471495472 | 4.46E-08 | peptidylprolyl cis/trans isomerase, NIMA-interacting 4 [Source:HGNC Symbol;Acc:HGNC:8992] |
| **ENSG00000116157** | GPX7 | 2882 | 198 | 49.45314476 | -1.175592452 | 0.260992656 | -4.504312379 | 6.66E-06 | glutathione peroxidase 7 [Source:HGNC Symbol;Acc:HGNC:4559] |
| **ENSG00000219665** | ZNF433-AS1 | NA | 197 | 76.16161272 | -1.076477976 | 0.197225705 | -5.458101815 | 4.81E-08 | ZNF433 and ZNF878 antisense RNA 1 [Source:HGNC Symbol;Acc:HGNC:53776] |
| **ENSG00000143416** | SELENBP1 | 8991 | 196 | 9.174928441 | -5.626291698 | 1.251351682 | -4.496171444 | 6.92E-06 | selenium binding protein 1 [Source:HGNC Symbol;Acc:HGNC:10719] |
| **ENSG00000178031** | ADAMTSL1 | 92949 | 195 | 59.12260065 | -7.11352469 | 0.862432513 | -8.248210242 | 1.61E-16 | ADAMTS like 1 [Source:HGNC Symbol;Acc:HGNC:14632] |
| **ENSG00000140678** | ITGAX | 3687 | 193 | 25.46423394 | -3.504827331 | 0.440154449 | -7.962721588 | 1.68E-15 | integrin subunit alpha X [Source:HGNC Symbol;Acc:HGNC:6152] |
| **ENSG00000160191** | PDE9A | 5152 | 192 | 285.1329577 | -1.886132674 | 0.136528631 | -13.8149241 | 2.07E-43 | phosphodiesterase 9A [Source:HGNC Symbol;Acc:HGNC:8795] |
| **ENSG00000099399** | MAGEB2 | 4113 | 191 | 52.5643312 | -3.151568888 | 0.301584642 | -10.4500311 | 1.46E-25 | MAGE family member B2 [Source:HGNC Symbol;Acc:HGNC:6809] |
| **ENSG00000022267** | FHL1 | 2273 | 190 | 489.7351177 | -1.269307148 | 0.115634867 | -10.97685485 | 4.94E-28 | four and a half LIM domains 1 [Source:HGNC Symbol;Acc:HGNC:3702] |
| **ENSG00000143507** | DUSP10 | 11221 | 189 | 258.4561519 | -0.699514715 | 0.109561133 | -6.384697681 | 1.72E-10 | dual specificity phosphatase 10 [Source:HGNC Symbol;Acc:HGNC:3065] |
| **ENSG00000109861** | CTSC | 1075 | 188 | 382.2021163 | -1.132987066 | 0.128797624 | -8.796645729 | 1.41E-18 | cathepsin C [Source:HGNC Symbol;Acc:HGNC:2528] |
| **ENSG00000142621** | FHAD1 | 114827 | 187 | 229.9370178 | -0.829532822 | 0.124404787 | -6.668013692 | 2.59E-11 | forkhead associated phosphopeptide binding domain 1 [Source:HGNC Symbol;Acc:HGNC:29408] |
| **ENSG00000183287** | CCBE1 | 147372 | 186 | 143.4809057 | -1.260743434 | 0.180973123 | -6.966467787 | 3.25E-12 | collagen and calcium binding EGF domains 1 [Source:HGNC Symbol;Acc:HGNC:29426] |
| **ENSG00000157601** | MX1 | 4599 | 185 | 204.2734451 | -2.397744503 | 0.191283062 | -12.53505917 | 4.8E-36 | MX dynamin like GTPase 1 [Source:HGNC Symbol;Acc:HGNC:7532] |
| **ENSG00000132535** | DLG4 | 1742 | 184 | 217.0253589 | -1.26497935 | 0.160048831 | -7.903708787 | 2.71E-15 | discs large MAGUK scaffold protein 4 [Source:HGNC Symbol;Acc:HGNC:2903] |
| **ENSG00000160298** | C21orf58 | 54058 | 183 | 190.484774 | -1.621936805 | 0.174106088 | -9.315796039 | 1.21E-20 | chromosome 21 open reading frame 58 [Source:HGNC Symbol;Acc:HGNC:1300] |
| **ENSG00000102048** | ASB9 | 140462 | 182 | 206.9285738 | -1.423313657 | 0.139558117 | -10.19871638 | 2.01E-24 | ankyrin repeat and SOCS box containing 9 [Source:HGNC Symbol;Acc:HGNC:17184] |
| **ENSG00000006210** | CX3CL1 | 6376 | 181 | 35.29575466 | -2.617326968 | 0.385066861 | -6.797071443 | 1.07E-11 | C-X3-C motif chemokine ligand 1 [Source:HGNC Symbol;Acc:HGNC:10647] |
| **ENSG00000108679** | LGALS3BP | 3959 | 180 | 706.0587837 | -2.090293226 | 0.15977226 | -13.08295462 | 4.12E-39 | galectin 3 binding protein [Source:HGNC Symbol;Acc:HGNC:6564] |
| **ENSG00000188783** | PRELP | 5549 | 179 | 19.08676785 | -4.068746444 | 0.58206965 | -6.990136732 | 2.75E-12 | proline and arginine rich end leucine rich repeat protein [Source:HGNC Symbol;Acc:HGNC:9357] |
| **ENSG00000173432** | SAA1 | 6288 | 178 | 78.11809159 | -2.758263738 | 0.238364289 | -11.57163158 | 5.74E-31 | serum amyloid A1 [Source:HGNC Symbol;Acc:HGNC:10513] |
| **ENSG00000175294** | CATSPER1 | 117144 | 177 | 65.6726391 | -1.763438603 | 0.24134308 | -7.306770952 | 2.74E-13 | cation channel sperm associated 1 [Source:HGNC Symbol;Acc:HGNC:17116] |
| **ENSG00000104419** | NDRG1 | 10397 | 176 | 258.5915067 | -1.468110999 | 0.156533278 | -9.378906633 | 6.67E-21 | N-myc downstream regulated 1 [Source:HGNC Symbol;Acc:HGNC:7679] |
| **ENSG00000156959** | LHFPL4 | 375323 | 175 | 92.52986166 | -1.507573635 | 0.229391036 | -6.57206864 | 4.96E-11 | LHFPL tetraspan subfamily member 4 [Source:HGNC Symbol;Acc:HGNC:29568] |
| **ENSG00000149557** | FEZ1 | 9638 | 174 | 79.86238906 | -2.180021408 | 0.217570843 | -10.01982333 | 1.25E-23 | fasciculation and elongation protein zeta 1 [Source:HGNC Symbol;Acc:HGNC:3659] |
| **ENSG00000174059** | CD34 | 947 | 173 | 76.94189246 | -1.578200028 | 0.206912845 | -7.627366146 | 2.4E-14 | CD34 molecule [Source:HGNC Symbol;Acc:HGNC:1662] |
| **ENSG00000183160** | TMEM119 | 338773 | 172 | 251.328967 | -3.194345813 | 0.296937093 | -10.75765167 | 5.45E-27 | transmembrane protein 119 [Source:HGNC Symbol;Acc:HGNC:27884] |
| **ENSG00000076706** | MCAM | 4162 | 171 | 1433.917112 | -2.387346936 | 0.136205413 | -17.52754816 | 8.83E-69 | melanoma cell adhesion molecule [Source:HGNC Symbol;Acc:HGNC:6934] |
| **ENSG00000001617** | SEMA3F | 6405 | 170 | 70.9042249 | -2.296808594 | 0.300952134 | -7.631806972 | 2.31E-14 | semaphorin 3F [Source:HGNC Symbol;Acc:HGNC:10728] |
| **ENSG00000169994** | MYO7B | 4648 | 169 | 468.6569745 | -1.457460149 | 0.160118109 | -9.102406675 | 8.84E-20 | myosin VIIB [Source:HGNC Symbol;Acc:HGNC:7607] |
| **ENSG00000058335** | RASGRF1 | 5923 | 168 | 29.76431607 | -3.708251296 | 0.387555624 | -9.56830727 | 1.09E-21 | Ras protein specific guanine nucleotide releasing factor 1 [Source:HGNC Symbol;Acc:HGNC:9875] |
| **ENSG00000171608** | PIK3CD | 5293 | 167 | 1291.971431 | -1.068394799 | 0.121689058 | -8.779711283 | 1.64E-18 | phosphatidylinositol-4,5-bisphosphate 3-kinase catalytic subunit delta [Source:HGNC Symbol;Acc:HGNC:8977] |
| **ENSG00000234380** | LINC01426 | NA | 166 | 37.30933079 | -3.453893089 | 0.38875642 | -8.884465715 | 6.42E-19 | long intergenic non-protein coding RNA 1426 [Source:HGNC Symbol;Acc:HGNC:50734] |
| **ENSG00000028137** | TNFRSF1B | 7133 | 165 | 100.4108151 | -2.989002991 | 0.269420375 | -11.09419802 | 1.34E-28 | TNF receptor superfamily member 1B [Source:HGNC Symbol;Acc:HGNC:11917] |
| **ENSG00000088882** | CPXM1 | 56265 | 164 | 268.2518574 | -2.278419746 | 0.184177661 | -12.37077142 | 3.76E-35 | carboxypeptidase X, M14 family member 1 [Source:HGNC Symbol;Acc:HGNC:15771] |
| **ENSG00000138964** | PARVG | 64098 | 163 | 8.432547075 | -5.008193835 | 1.096203784 | -4.56867045 | 4.91E-06 | parvin gamma [Source:HGNC Symbol;Acc:HGNC:14654] |
| **ENSG00000030304** | MUSK | 4593 | 162 | 40.66025609 | -1.608307237 | 0.243759297 | -6.597931888 | 4.17E-11 | muscle associated receptor tyrosine kinase [Source:HGNC Symbol;Acc:HGNC:7525] |
| **ENSG00000188064** | WNT7B | 7477 | 161 | 283.4068655 | -1.903170809 | 0.198404032 | -9.592399852 | 8.61E-22 | Wnt family member 7B [Source:HGNC Symbol;Acc:HGNC:12787] |
| **ENSG00000100626** | GALNT16 | 57452 | 160 | 92.03483689 | -1.820202684 | 0.196857056 | -9.246316709 | 2.32E-20 | polypeptide N-acetylgalactosaminyltransferase 16 [Source:HGNC Symbol;Acc:HGNC:23233] |
| **ENSG00000100342** | APOL1 | 8542 | 159 | 259.7144456 | -1.083959747 | 0.156535791 | -6.924676713 | 4.37E-12 | apolipoprotein L1 [Source:HGNC Symbol;Acc:HGNC:618] |
| **ENSG00000204103** | MAFB | 9935 | 158 | 16.98626028 | -2.077646575 | 0.411479789 | -5.049206866 | 4.44E-07 | MAF bZIP transcription factor B [Source:HGNC Symbol;Acc:HGNC:6408] |
| **ENSG00000134516** | DOCK2 | 1794 | 157 | 82.6388376 | -1.012308456 | 0.152985391 | -6.617026977 | 3.66E-11 | dedicator of cytokinesis 2 [Source:HGNC Symbol;Acc:HGNC:2988] |
| **ENSG00000135424** | ITGA7 | 3679 | 156 | 143.3263873 | -1.449222037 | 0.18298235 | -7.920009965 | 2.37E-15 | integrin subunit alpha 7 [Source:HGNC Symbol;Acc:HGNC:6143] |
| **ENSG00000007237** | GAS7 | 8522 | 155 | 340.4038808 | -1.417037332 | 0.126081721 | -11.23903859 | 2.62E-29 | growth arrest specific 7 [Source:HGNC Symbol;Acc:HGNC:4169] |
| **ENSG00000138395** | CDK15 | 65061 | 154 | 44.75306801 | -4.345568655 | 0.442805352 | -9.813722059 | 9.83E-23 | cyclin dependent kinase 15 [Source:HGNC Symbol;Acc:HGNC:14434] |
| **ENSG00000165071** | TMEM71 | 137835 | 153 | 3.711185495 | -5.916570383 | 1.291968706 | -4.579499763 | 4.66E-06 | transmembrane protein 71 [Source:HGNC Symbol;Acc:HGNC:26572] |
| **ENSG00000165959** | CLMN | 79789 | 152 | 88.26540724 | -1.681952596 | 0.176183093 | -9.546617509 | 1.34E-21 | calmin [Source:HGNC Symbol;Acc:HGNC:19972] |
| **ENSG00000178538** | CA8 | 767 | 151 | 240.2527914 | -3.397358874 | 0.154508548 | -21.9881613 | 3.74E-107 | carbonic anhydrase 8 [Source:HGNC Symbol;Acc:HGNC:1382] |
| **ENSG00000261761** | LINC02616 | NA | 150 | 7.11447832 | -7.614418372 | 1.302292991 | -5.846931855 | 5.01E-09 | long intergenic non-protein coding RNA 2616 [Source:HGNC Symbol;Acc:HGNC:54078] |
| **ENSG00000160219** | GAB3 | 139716 | 149 | 35.14442464 | -1.931096649 | 0.22712901 | -8.502201659 | 1.86E-17 | GRB2 associated binding protein 3 [Source:HGNC Symbol;Acc:HGNC:17515] |
| **ENSG00000041982** | TNC | 3371 | 148 | 1038.988883 | -1.192705673 | 0.179708131 | -6.636904329 | 3.2E-11 | tenascin C [Source:HGNC Symbol;Acc:HGNC:5318] |
| **ENSG00000198585** | NUDT16 | 131870 | 147 | 268.064243 | -0.911354014 | 0.122830185 | -7.419625844 | 1.17E-13 | nudix hydrolase 16 [Source:HGNC Symbol;Acc:HGNC:26442] |
| **ENSG00000084234** | APLP2 | 334 | 146 | 1317.181898 | -1.240670038 | 0.142871989 | -8.683787828 | 3.83E-18 | amyloid beta precursor like protein 2 [Source:HGNC Symbol;Acc:HGNC:598] |
| **ENSG00000130702** | LAMA5 | 3911 | 145 | 814.5235712 | -0.633889969 | 0.149450432 | -4.241472987 | 2.22E-05 | laminin subunit alpha 5 [Source:HGNC Symbol;Acc:HGNC:6485] |
| **ENSG00000137203** | TFAP2A | 7020 | 144 | 173.0363684 | -1.184576862 | 0.129371472 | -9.156399368 | 5.37E-20 | transcription factor AP-2 alpha [Source:HGNC Symbol;Acc:HGNC:11742] |
| **ENSG00000146950** | SHROOM2 | 357 | 142 | 294.1338521 | -0.935671649 | 0.134189404 | -6.972768511 | 3.11E-12 | shroom family member 2 [Source:HGNC Symbol;Acc:HGNC:630] |
| **ENSG00000138798** | EGF | 1950 | 141 | 14.66570779 | -3.725145139 | 0.515415048 | -7.22746678 | 4.92E-13 | epidermal growth factor [Source:HGNC Symbol;Acc:HGNC:3229] |
| **ENSG00000165124** | SVEP1 | 79987 | 140 | 42.65244567 | -1.97576714 | 0.238205148 | -8.29439313 | 1.09E-16 | sushi, von Willebrand factor type A, EGF and pentraxin domain containing 1 [Source:HGNC Symbol;Acc:HGNC:1 |
| **ENSG00000050767** | COL23A1 | 91522 | 139 | 59.82289889 | -1.905216862 | 0.239451744 | -7.956579591 | 1.77E-15 | collagen type XXIII alpha 1 chain [Source:HGNC Symbol;Acc:HGNC:22990] |
| **ENSG00000133195** | SLC39A11 | 201266 | 138 | 333.0534858 | -0.699805759 | 0.10162273 | -6.886311322 | 5.73E-12 | solute carrier family 39 member 11 [Source:HGNC Symbol;Acc:HGNC:14463] |
| **ENSG00000189143** | CLDN4 | 1364 | 137 | 102.7408441 | -1.154944526 | 0.189845111 | -6.0836148 | 1.18E-09 | claudin 4 [Source:HGNC Symbol;Acc:HGNC:2046] |
| **ENSG00000181264** | TLCD5 | 219902 | 136 | 111.5915889 | -1.017560765 | 0.151835472 | -6.701732818 | 2.06E-11 | TLC domain containing 5 [Source:HGNC Symbol;Acc:HGNC:28280] |
| **ENSG00000213654** | GPSM3 | 63940 | 135 | 228.1032992 | -1.122640731 | 0.207099034 | -5.420791729 | 5.93E-08 | G protein signaling modulator 3 [Source:HGNC Symbol;Acc:HGNC:13945] |
| **ENSG00000182492** | BGN | 633 | 134 | 1221.216004 | -1.324471335 | 0.1897822 | -6.978901784 | 2.97E-12 | biglycan [Source:HGNC Symbol;Acc:HGNC:1044] |
| **ENSG00000142173** | COL6A2 | 1292 | 133 | 6948.413222 | -1.076581646 | 0.168634493 | -6.384112922 | 1.72E-10 | collagen type VI alpha 2 chain [Source:HGNC Symbol;Acc:HGNC:2212] |
| **ENSG00000186470** | BTN3A2 | 11118 | 132 | 205.5227998 | -1.220302487 | 0.144327422 | -8.455097962 | 2.79E-17 | butyrophilin subfamily 3 member A2 [Source:HGNC Symbol;Acc:HGNC:1139] |
| **ENSG00000163703** | CRELD1 | 78987 | 131 | 263.6915069 | -1.014731317 | 0.176735127 | -5.741537268 | 9.38E-09 | cysteine rich with EGF like domains 1 [Source:HGNC Symbol;Acc:HGNC:14630] |
| **ENSG00000196460** | RFX8 | 731220 | 130 | 17.88140664 | -2.545766339 | 0.575124235 | -4.426463335 | 9.58E-06 | RFX family member 8, lacking RFX DNA binding domain [Source:HGNC Symbol;Acc:HGNC:37253] |
| **ENSG00000115457** | IGFBP2 | 3485 | 128 | 934.7831821 | -0.924911192 | 0.219161635 | -4.220223995 | 2.44E-05 | insulin like growth factor binding protein 2 [Source:HGNC Symbol;Acc:HGNC:5471] |

| | | | | | | | | | |
|---|---|---|---|---|---|---|---|---|---|
| ENSG00000162511 | LAPTM5 | 7805 | 127 | 61.9029448 | -1.950137203 | 0.241000072 | -8.091853195 | 5.88E-16 | lysosomal protein transmembrane 5 [Source:HGNC Symbol;Acc:HGNC:29612] |
| ENSG00000160255 | ITGB2 | 3689 | 126 | 171.7893614 | -1.18962744 | 0.159796576 | -7.444636598 | 9.72E-14 | integrin subunit beta 2 [Source:HGNC Symbol;Acc:HGNC:6155] |
| ENSG00000157303 | SUSD3 | 203328 | 125 | 170.8680292 | -0.742091788 | 0.197685719 | -3.753896797 | 0.000174107 | sushi domain containing 3 [Source:HGNC Symbol;Acc:HGNC:28391] |
| ENSG00000105929 | ATP6V0A4 | 50617 | 124 | 29.57627406 | -0.762903563 | 0.238381002 | -3.20035387 | 0.00137259 | ATPase H+ transporting V0 subunit a4 [Source:HGNC Symbol;Acc:HGNC:866] |
| ENSG00000204291 | COL15A1 | 1306 | 123 | 44.19679194 | -1.215527413 | 0.301587906 | -4.030424921 | 5.57E-05 | collagen type XV alpha 1 chain [Source:HGNC Symbol;Acc:HGNC:2192] |
| ENSG00000185686 | PRAME | 23532 | 122 | 360.8696045 | -0.890076266 | 0.127313523 | -6.991215423 | 2.73E-12 | preferentially expressed antigen in melanoma [Source:HGNC Symbol;Acc:HGNC:9336] |
| ENSG00000144115 | THNSL2 | 55258 | 121 | 21.44919615 | -1.030303327 | 0.347052363 | -2.968725865 | 0.002990372 | threonine synthase like 2 [Source:HGNC Symbol;Acc:HGNC:25602] |
| ENSG00000067445 | TRO | 7216 | 120 | 217.9025489 | -0.78170635 | 0.152154428 | -5.137585283 | 2.78E-07 | trophinin [Source:HGNC Symbol;Acc:HGNC:12326] |
| ENSG00000177706 | FAM20C | 56975 | 119 | 2663.376615 | -0.793906688 | 0.146293653 | -5.426801999 | 5.74E-08 | FAM20C golgi associated secretory pathway kinase [Source:HGNC Symbol;Acc:HGNC:22140] |
| ENSG00000123453 | SARDH | 1757 | 118 | 90.24681971 | -0.732595086 | 0.187232217 | -3.912761909 | 9.12E-05 | sarcosine dehydrogenase [Source:HGNC Symbol;Acc:HGNC:10536] |
| ENSG00000120937 | NPPB | 4879 | 117 | 7.452570779 | -1.191961402 | 0.598997449 | -1.989927344 | 0.04659894 | natriuretic peptide B [Source:HGNC Symbol;Acc:HGNC:7940] |
| ENSG00000083750 | RRAGB | 10325 | 116 | 35.22183348 | -1.110096424 | 0.226640139 | -4.898057455 | 9.68E-07 | Ras related GTP binding B [Source:HGNC Symbol;Acc:HGNC:19901] |
| ENSG00000147202 | DIAPH2 | 1730 | 115 | 6.334784452 | -3.486423372 | 0.722829321 | -4.823300982 | 1.41E-06 | diaphanous related formin 2 [Source:HGNC Symbol;Acc:HGNC:2877] |
| ENSG00000213759 | UGT2B11 | 10720 | 114 | 17.21461076 | -3.059284201 | 0.446873543 | -6.845972988 | 7.6E-12 | UDP glucuronosyltransferase family 2 member B11 [Source:HGNC Symbol;Acc:HGNC:12545] |
| ENSG00000224184 | MIR3681HG | NA | 112 | 9.590563 | -3.151834932 | 0.584327861 | -5.393949431 | 6.89E-08 | MIR3681 host gene [Source:HGNC Symbol;Acc:HGNC:52001] |
| ENSG00000065618 | COL17A1 | 1308 | 111 | 18.51926349 | -3.895064863 | 0.600335411 | -6.488147776 | 8.69E-11 | collagen type XVII alpha 1 chain [Source:HGNC Symbol;Acc:HGNC:2194] |
| ENSG00000151702 | FLI1 | 2313 | 110 | 132.8982471 | -1.340029211 | 0.156285972 | -8.574212976 | 9.98E-18 | Fli-1 proto-oncogene, ETS transcription factor [Source:HGNC Symbol;Acc:HGNC:3749] |
| ENSG00000112297 | CRYBG1 | 202 | 109 | 123.2291647 | -1.587289858 | 0.196718603 | -8.068834551 | 7.1E-16 | crystallin beta-gamma domain containing 1 [Source:HGNC Symbol;Acc:HGNC:356] |
| ENSG00000188158 | NHS | 4810 | 108 | 9.986941469 | -2.975375438 | 0.493659765 | -6.027178328 | 1.67E-09 | NHS actin remodeling regulator [Source:HGNC Symbol;Acc:HGNC:7820] |
| ENSG00000145730 | PAM | 5066 | 107 | 366.8405667 | -0.831921034 | 0.141785063 | -5.867480079 | 4.42E-09 | peptidylglycine alpha-amidating monooxygenase [Source:HGNC Symbol;Acc:HGNC:8596] |
| ENSG00000078124 | ACER3 | 55331 | 105 | 211.0907109 | -0.850168497 | 0.155452647 | -5.468986961 | 4.53E-08 | alkaline ceramidase 3 [Source:HGNC Symbol;Acc:HGNC:16066] |
| ENSG00000196177 | ACADSB | 36 | 104 | 64.93959837 | -1.184906488 | 0.219638925 | -5.394792796 | 6.86E-08 | acyl-CoA dehydrogenase short/branched chain [Source:HGNC Symbol;Acc:HGNC:91] |
| ENSG00000261371 | PECAM1 | 5175 | 103 | 17.91889676 | -4.296645414 | 0.644477895 | -6.66686235 | 2.61E-11 | platelet and endothelial cell adhesion molecule 1 [Source:HGNC Symbol;Acc:HGNC:8823] |
| ENSG00000137463 | MGARP | 84709 | 102 | 43.01939235 | -0.960617798 | 0.239284497 | -4.014542558 | 5.96E-05 | mitochondria localized glutamic acid rich protein [Source:HGNC Symbol;Acc:HGNC:29969] |
| ENSG00000124785 | NRN1 | 51299 | 101 | 73.68785386 | -2.520073462 | 0.250848488 | -10.04619753 | 9.55E-24 | neuritin 1 [Source:HGNC Symbol;Acc:HGNC:17972] |
| ENSG00000181467 | RAP2B | 5912 | 100 | 134.8004641 | -0.621464118 | 0.137636618 | -4.515252757 | 6.32E-06 | RAP2B, member of RAS oncogene family [Source:HGNC Symbol;Acc:HGNC:9862] |
| ENSG00000144583 | MARCHF4 | 57574 | 99 | 39.40796104 | -1.765889089 | 0.248058681 | -7.118836069 | 1.09E-12 | membrane associated ring-CH-type finger 4 [Source:HGNC Symbol;Acc:HGNC:29269] |
| ENSG00000158458 | NRG2 | 9542 | 98 | 31.70234636 | -2.857348394 | 0.353331685 | -8.086872793 | 6.12E-16 | neuregulin 2 [Source:HGNC Symbol;Acc:HGNC:7998] |
| ENSG00000128284 | APOL3 | 80833 | 97 | 58.37850508 | -2.315042472 | 0.214198791 | -10.80791571 | 3.16E-27 | apolipoprotein L3 [Source:HGNC Symbol;Acc:HGNC:14868] |
| ENSG00000198753 | PLXNB3 | 5365 | 96 | 35.6845025 | -1.718167841 | 0.316324602 | -5.431660486 | 5.58E-08 | plexin B3 [Source:HGNC Symbol;Acc:HGNC:9105] |
| ENSG00000258183 | LINC02392 | NA | 95 | 59.10712565 | -2.671478275 | 0.252078107 | -10.59781949 | 3.05E-26 | long intergenic non-protein coding RNA 2392 [Source:HGNC Symbol;Acc:HGNC:53319] |
| ENSG00000049449 | RCN1 | 5954 | 94 | 125.5211653 | -1.997813405 | 0.173926449 | -11.48654171 | 1.54E-30 | reticulocalbin 1 [Source:HGNC Symbol;Acc:HGNC:9934] |
| ENSG00000079263 | SP140 | 11262 | 93 | 68.51547527 | -2.614313886 | 0.232018918 | -11.26767553 | 1.9E-29 | SP140 nuclear body protein [Source:HGNC Symbol;Acc:HGNC:17133] |
| ENSG00000122378 | PRXL2A | 84293 | 92 | 71.52115588 | -2.232623729 | 0.198772663 | -11.23204615 | 2.84E-29 | peroxiredoxin like 2A [Source:HGNC Symbol;Acc:HGNC:28651] |
| ENSG00000175093 | SPSB4 | 92369 | 91 | 22.25460831 | -1.75427003 | 0.32007485 | -5.480811846 | 4.23E-08 | splA/ryanodine receptor domain and SOCS box containing 4 [Source:HGNC Symbol;Acc:HGNC:30630] |
| ENSG00000185201 | IFITM2 | 10581 | 90 | 130.7381763 | -1.982128502 | 0.200943216 | -9.864122537 | 5.96E-23 | interferon induced transmembrane protein 2 [Source:HGNC Symbol;Acc:HGNC:5413] |
| ENSG00000149292 | TTC12 | 54970 | 89 | 32.49033132 | -2.413649476 | 0.292115075 | -8.262666615 | 1.42E-16 | tetratricopeptide repeat domain 12 [Source:HGNC Symbol;Acc:HGNC:23700] |
| ENSG00000106868 | SUSD1 | 64420 | 88 | 127.4509845 | -0.742722749 | 0.135776961 | -5.470167728 | 4.5E-08 | sushi domain containing 1 [Source:HGNC Symbol;Acc:HGNC:25413] |
| ENSG00000162441 | LZIC | 84328 | 87 | 301.2743977 | -0.850020682 | 0.114821215 | -7.402993292 | 1.33E-13 | leucine zipper and CTNNBIP1 domain containing [Source:HGNC Symbol;Acc:HGNC:17497] |
| ENSG00000196639 | HRH1 | 3269 | 86 | 350.7500031 | -0.760981988 | 0.094393258 | -8.061825619 | 7.52E-16 | histamine receptor H1 [Source:HGNC Symbol;Acc:HGNC:5182] |
| ENSG00000135318 | NT5E | 4907 | 85 | 377.2716698 | -1.51211786 | 0.122943193 | -12.29932156 | 9.13E-35 | 5'-nucleotidase ecto [Source:HGNC Symbol;Acc:HGNC:8021] |
| ENSG00000214548 | MEG3 | NA | 84 | 14.0777437 | -4.338749702 | 0.578124896 | -7.504865694 | 6.15E-14 | maternally expressed 3 [Source:HGNC Symbol;Acc:HGNC:14575] |
| ENSG00000138496 | PARP9 | 83666 | 83 | 74.46171713 | -1.224406172 | 0.183572431 | -6.669880462 | 2.56E-11 | poly(ADP-ribose) polymerase family member 9 [Source:HGNC Symbol;Acc:HGNC:24118] |
| ENSG00000196730 | DAPK1 | 1612 | 82 | 42.73672522 | -1.841939181 | 0.236455287 | -7.789799093 | 6.71E-15 | death associated protein kinase 1 [Source:HGNC Symbol;Acc:HGNC:2674] |
| ENSG00000162433 | AK4 | 205 | 81 | 117.9357781 | -1.204263949 | 0.13927529 | -8.646644688 | 5.3E-18 | adenylate kinase 4 [Source:HGNC Symbol;Acc:HGNC:363] |
| ENSG00000169247 | SH3TC2 | 79628 | 80 | 170.5441139 | -0.741918656 | 0.119332604 | -6.217233455 | 5.06E-10 | SH3 domain and tetratricopeptide repeats 2 [Source:HGNC Symbol;Acc:HGNC:29427] |
| ENSG00000159167 | STC1 | 6781 | 79 | 89.50391616 | -1.271845747 | 0.167602899 | -7.588447186 | 3.24E-14 | stanniocalcin 1 [Source:HGNC Symbol;Acc:HGNC:11373] |
| ENSG00000158270 | COLEC12 | 81035 | 78 | 334.0143966 | -0.983424202 | 0.142767813 | -6.88827673 | 5.65E-12 | collectin subfamily member 12 [Source:HGNC Symbol;Acc:HGNC:16016] |
| ENSG00000177675 | CD163L1 | 283316 | 77 | 72.29687436 | -1.321107213 | 0.241950459 | -5.460238511 | 4.75E-08 | CD163 molecule like 1 [Source:HGNC Symbol;Acc:HGNC:30375] |
| ENSG00000064309 | CDON | 50937 | 76 | 34.61523189 | -1.981680266 | 0.268799476 | -7.37233679 | 1.68E-13 | cell adhesion associated, oncogene regulated [Source:HGNC Symbol;Acc:HGNC:17104] |
| ENSG00000147862 | NFIB | 4781 | 75 | 122.5089365 | -1.326094856 | 0.206323419 | -6.427262892 | 1.3E-10 | nuclear factor I B [Source:HGNC Symbol;Acc:HGNC:7785] |
| ENSG00000065534 | MYLK | 4638 | 74 | 221.7420353 | -1.403273906 | 0.148504262 | -9.44938472 | 3.41E-21 | myosin light chain kinase [Source:HGNC Symbol;Acc:HGNC:7590] |
| ENSG00000146938 | NLGN4X | 57502 | 73 | 282.0650059 | -1.531307973 | 0.167649885 | -9.133963744 | 6.6E-20 | neuroligin 4 X-linked [Source:HGNC Symbol;Acc:HGNC:14287] |
| ENSG00000154556 | SORBS2 | 8470 | 72 | 155.0597443 | -1.470338932 | 0.180507533 | -8.14558211 | 3.77E-16 | sorbin and SH3 domain containing 2 [Source:HGNC Symbol;Acc:HGNC:24098] |
| ENSG00000033122 | LRRC7 | 57554 | 71 | 23.49335009 | -2.110920833 | 0.304894317 | -6.923450899 | 4.41E-12 | leucine rich repeat containing 7 [Source:HGNC Symbol;Acc:HGNC:18531] |
| ENSG00000169306 | IL1RAPL1 | 11141 | 70 | 44.05084833 | -2.253377652 | 0.336485175 | -6.696811083 | 2.13E-11 | interleukin 1 receptor accessory protein like 1 [Source:HGNC Symbol;Acc:HGNC:5996] |
| ENSG00000096060 | FKBP5 | 2289 | 68 | 88.6937815 | -1.366574465 | 0.170877257 | -7.997404044 | 1.27E-15 | FKBP prolyl isomerase 5 [Source:HGNC Symbol;Acc:HGNC:3721] |
| ENSG00000145632 | PLK2 | 10769 | 67 | 603.0619635 | -1.267510521 | 0.127423896 | -9.947196394 | 2.59E-23 | polo like kinase 2 [Source:HGNC Symbol;Acc:HGNC:19699] |
| ENSG00000153767 | GTF2E1 | 2960 | 66 | 75.83249082 | -1.080448487 | 0.191896583 | -5.630368562 | 1.8E-08 | general transcription factor IIE subunit 1 [Source:HGNC Symbol;Acc:HGNC:4650] |
| ENSG00000050344 | NFE2L3 | 9603 | 65 | 166.2742673 | -1.551232282 | 0.177261207 | -8.751109763 | 2.11E-18 | nuclear factor, erythroid 2 like 3 [Source:HGNC Symbol;Acc:HGNC:7783] |
| ENSG00000133393 | FOPNL | 123811 | 64 | 246.2025805 | -0.870954668 | 0.129058573 | -6.748522384 | 1.49E-11 | FGFR1OP N-terminal like [Source:HGNC Symbol;Acc:HGNC:26435] |
| ENSG00000152661 | GJA1 | 2697 | 63 | 133.0112079 | -2.391409444 | 0.1956995 | -12.21980357 | 2.44E-34 | gap junction protein alpha 1 [Source:HGNC Symbol;Acc:HGNC:4274] |
| ENSG00000169083 | AR | 367 | 62 | 34.7777221 | -1.710428879 | 0.235735825 | -7.255701935 | 4E-13 | androgen receptor [Source:HGNC Symbol;Acc:HGNC:644] |
| ENSG00000198121 | LPAR1 | 1902 | 61 | 334.5648867 | -1.161158062 | 0.161865688 | -7.173589901 | 7.31E-13 | lysophosphatidic acid receptor 1 [Source:HGNC Symbol;Acc:HGNC:3166] |
| ENSG00000106049 | HIBADH | 11112 | 60 | 159.5481836 | -1.389478158 | 0.206032926 | -6.743961687 | 1.54E-11 | 3-hydroxyisobutyrate dehydrogenase [Source:HGNC Symbol;Acc:HGNC:4907] |
| ENSG00000138759 | FRAS1 | 80144 | 59 | 41.01780323 | -1.689431278 | 0.215681373 | -7.832995746 | 4.76E-15 | Fraser extracellular matrix complex subunit 1 [Source:HGNC Symbol;Acc:HGNC:19185] |
| ENSG00000169446 | MMGT1 | 93380 | 58 | 369.781027 | -1.065437166 | 0.109861408 | -9.698011246 | 3.07E-22 | membrane magnesium transporter 1 [Source:HGNC Symbol;Acc:HGNC:28100] |
| ENSG00000114166 | KAT2B | 8850 | 57 | 67.87638451 | -1.823571368 | 0.258892676 | -7.043734876 | 1.87E-12 | lysine acetyltransferase 2B [Source:HGNC Symbol;Acc:HGNC:8638] |
| ENSG00000109819 | PPARGC1A | 10891 | 56 | 2.969906582 | -3.704636152 | 0.979555434 | -3.781956612 | 0.0001556 | PPARG coactivator 1 alpha [Source:HGNC Symbol;Acc:HGNC:9237] |
| ENSG00000197372 | ZNF675 | 171392 | 55 | 335.6105243 | -0.881458573 | 0.123022598 | -7.165013465 | 7.78E-13 | zinc finger protein 675 [Source:HGNC Symbol;Acc:HGNC:30768] |
| ENSG00000005893 | LAMP2 | 3920 | 54 | 1437.33728 | -0.685113201 | 0.096758943 | -7.080618873 | 1.44E-12 | lysosomal associated membrane protein 2 [Source:HGNC Symbol;Acc:HGNC:6501] |
| ENSG00000111846 | GCNT2 | 2651 | 53 | 166.9527499 | -1.254180395 | 0.165713395 | -7.568370646 | 3.78E-14 | glucosaminyl (N-acetyl) transferase 2 (I blood group) [Source:HGNC Symbol;Acc:HGNC:4204] |
| ENSG00000144959 | NCEH1 | 57552 | 52 | 131.4529735 | -0.957784539 | 0.175158834 | -5.468091545 | 4.55E-08 | neutral cholesterol ester hydrolase 1 [Source:HGNC Symbol;Acc:HGNC:29260] |
| ENSG00000157766 | ACAN | 176 | 51 | 11.52482147 | -3.081214812 | 0.515827416 | -5.973344407 | 2.32E-09 | aggrecan [Source:HGNC Symbol;Acc:HGNC:319] |
| ENSG00000161896 | IP6K3 | 117283 | 50 | 26.04327393 | -1.393116253 | 0.296635591 | -4.696389424 | 2.65E-06 | inositol hexakisphosphate kinase 3 [Source:HGNC Symbol;Acc:HGNC:17269] |
| ENSG00000160282 | FTCD | 10841 | 49 | 14.42522191 | -2.32876975 | 0.427712141 | -5.444712752 | 5.19E-08 | formimidoyltransferase cyclodeaminase [Source:HGNC Symbol;Acc:HGNC:3974] |
| ENSG00000143786 | CNIH3 | 149111 | 48 | 44.62630891 | -1.778851312 | 0.256477289 | -6.935706926 | 4.04E-12 | cornichon family AMPA receptor auxiliary protein 3 [Source:HGNC Symbol;Acc:HGNC:26802] |
| ENSG00000196220 | SRGAP3 | 9901 | 47 | 20.12754286 | -2.019046283 | 0.317494092 | -6.359319235 | 2.03E-10 | SLIT-ROBO Rho GTPase activating protein 3 [Source:HGNC Symbol;Acc:HGNC:19744] |
| ENSG00000144681 | STAC | 6769 | 45 | 34.08169804 | -1.752973788 | 0.277241269 | -6.322917928 | 2.57E-10 | SH3 and cysteine rich domain [Source:HGNC Symbol;Acc:HGNC:11353] |
| ENSG00000166426 | CRABP1 | 1381 | 44 | 9.151373705 | -5.89671295 | 1.245047428 | -4.736135199 | 2.18E-06 | cellular retinoic acid binding protein 1 [Source:HGNC Symbol;Acc:HGNC:2338] |
| ENSG00000189337 | KAZN | 23254 | 43 | 189.1084341 | -0.615342518 | 0.118444767 | -5.195185359 | 2.05E-07 | kazrin, periplakin interacting protein [Source:HGNC Symbol;Acc:HGNC:29173] |
| ENSG00000234323 | LINC01505 | NA | 42 | 9.718072006 | -3.230669508 | 0.593982879 | -5.438994325 | 5.36E-08 | long intergenic non-protein coding RNA 1505 [Source:HGNC Symbol;Acc:HGNC:51186] |
| ENSG00000154175 | ABI3BP | 25890 | 41 | 33.28890654 | -1.492664478 | 0.308238627 | -4.842561403 | 1.28E-06 | ABI family member 3 binding protein [Source:HGNC Symbol;Acc:HGNC:17265] |
| ENSG00000006788 | MYH13 | 8735 | 40 | 2.868562721 | -6.326614776 | 1.397796135 | -4.526135549 | 6.01E-06 | myosin heavy chain 13 [Source:HGNC Symbol;Acc:HGNC:7571] |
| ENSG00000186684 | CYP27C1 | 339761 | 39 | 26.97659291 | -0.882714795 | 0.355851447 | -2.480571044 | 0.013117212 | cytochrome P450 family 27 subfamily C member 1 [Source:HGNC Symbol;Acc:HGNC:33480] |
| ENSG00000091409 | ITGA6 | 3655 | 38 | 35.32527278 | -1.93492003 | 0.270153461 | -7.16229961 | 7.93E-13 | integrin subunit alpha 6 [Source:HGNC Symbol;Acc:HGNC:6142] |

| | | | | | | | | | |
|---|---|---|---|---|---|---|---|---|---|
| **ENSG00000164684** | ZNF704 | 619279 | 37 | 57.36866785 | -1.057112099 | 0.21435784 | -4.931529909 | 8.16E-07 | zinc finger protein 704 [Source:HGNC Symbol;Acc:HGNC:32291] |
| **ENSG00000001561** | ENPP4 | 22875 | 36 | 12.20646505 | -1.965304001 | 0.416323115 | -4.720621874 | 2.35E-06 | ectonucleotide pyrophosphatase/phosphodiesterase 4 [Source:HGNC Symbol;Acc:HGNC:3359] |
| **ENSG00000178202** | POGLUT3 | 143888 | 35 | 338.6668492 | -1.369455071 | 0.25273212 | -5.418603199 | 6.01E-08 | protein O-glucosyltransferase 3 [Source:HGNC Symbol;Acc:HGNC:28496] |
| **ENSG00000171867** | PRNP | 5621 | 34 | 739.6520221 | -0.869920406 | 0.132918116 | -6.544784336 | 5.96E-11 | prion protein [Source:HGNC Symbol;Acc:HGNC:9449] |
| **ENSG00000140367** | UBE2Q2 | 92912 | 33 | 150.7054772 | -1.407005843 | 0.245117856 | -5.740119725 | 9.46E-09 | ubiquitin conjugating enzyme E2 Q2 [Source:HGNC Symbol;Acc:HGNC:19248] |
| **ENSG00000082701** | GSK3B | 2932 | 32 | 443.5888393 | -0.59747232 | 0.14093613 | -4.239312658 | 2.24E-05 | glycogen synthase kinase 3 beta [Source:HGNC Symbol;Acc:HGNC:4617] |
| **ENSG00000101871** | MID1 | 4281 | 31 | 1020.492372 | -0.994334554 | 0.148837141 | -6.680688339 | 2.38E-11 | midline 1 [Source:HGNC Symbol;Acc:HGNC:7095] |
| **ENSG00000133835** | HSD17B4 | 3295 | 30 | 326.0065071 | -0.677877201 | 0.138869586 | -4.881394268 | 1.05E-06 | hydroxysteroid 17-beta dehydrogenase 4 [Source:HGNC Symbol;Acc:HGNC:5213] |
| **ENSG00000145819** | ARHGAP26 | 23092 | 29 | 151.9507233 | -0.815464624 | 0.16721004 | -4.876887918 | 1.08E-06 | Rho GTPase activating protein 26 [Source:HGNC Symbol;Acc:HGNC:17073] |
| **ENSG00000027697** | IFNGR1 | 3459 | 28 | 189.9437664 | -0.920058208 | 0.170881308 | -5.384194561 | 7.28E-08 | interferon gamma receptor 1 [Source:HGNC Symbol;Acc:HGNC:5439] |
| **ENSG00000196937** | FAM3C | 10447 | 27 | 213.4378552 | -1.218134078 | 0.169316792 | -7.194407978 | 6.27E-13 | FAM3 metabolism regulating signaling molecule C [Source:HGNC Symbol;Acc:HGNC:18664] |
| **ENSG00000112186** | CAP2 | 10486 | 26 | 381.978783 | -0.75506647 | 0.130216337 | -5.798554063 | 6.69E-09 | cyclase associated actin cytoskeleton regulatory protein 2 [Source:HGNC Symbol;Acc:HGNC:20039] |
| **ENSG00000174844** | DNAH12 | 201625 | 25 | 26.81751841 | -2.234096354 | 0.39372744 | -5.674220603 | 1.39E-08 | dynein axonemal heavy chain 12 [Source:HGNC Symbol;Acc:HGNC:2943] |
| **ENSG00000163395** | IGFN1 | 91156 | 24 | 111.819379 | -1.479215786 | 0.221479721 | -6.678786564 | 2.41E-11 | immunoglobulin like and fibronectin type III domain containing 1 [Source:HGNC Symbol;Acc:HGNC:24607] |
| **ENSG00000138134** | STAMBPL1 | 57559 | 23 | 49.14379247 | -1.182771447 | 0.258459659 | -4.576232323 | 4.73E-06 | STAM binding protein like 1 [Source:HGNC Symbol;Acc:HGNC:24105] |
| **ENSG00000185009** | AP3M1 | 26985 | 22 | 191.4552484 | -1.149119484 | 0.243937399 | -4.710714665 | 2.47E-06 | adaptor related protein complex 3 subunit mu 1 [Source:HGNC Symbol;Acc:HGNC:569] |
| **ENSG00000023608** | SNAPC1 | 6617 | 21 | 57.06277954 | -0.665667927 | 0.299753131 | -2.220720511 | 0.026369897 | small nuclear RNA activating complex polypeptide 1 [Source:HGNC Symbol;Acc:HGNC:11134] |
| **ENSG00000170854** | RIOX2 | 84864 | 20 | 129.1073237 | -0.806889276 | 0.199116974 | -4.052337991 | 5.07E-05 | ribosomal oxygenase 2 [Source:HGNC Symbol;Acc:HGNC:19441] |
| **ENSG00000148516** | ZEB1 | 6935 | 19 | 69.25908653 | -1.489476472 | 0.342373474 | -4.350443554 | 1.36E-05 | zinc finger E-box binding homeobox 1 [Source:HGNC Symbol;Acc:HGNC:11642] |
| **ENSG00000189180** | ZNF33A | 7581 | 18 | 88.8758097 | -1.315869474 | 0.242556866 | -5.424993707 | 5.8E-08 | zinc finger protein 33A [Source:HGNC Symbol;Acc:HGNC:13096] |
| **ENSG00000087095** | NLK | 51701 | 17 | 90.14039085 | -0.947139304 | 0.19203565 | -4.932101443 | 8.13E-07 | nemo like kinase [Source:HGNC Symbol;Acc:HGNC:29858] |
| **ENSG00000175175** | PPM1E | 22843 | 16 | 43.4090471 | -0.755109362 | 0.234568852 | -3.21913739 | 0.001285769 | protein phosphatase, Mg2+/Mn2+ dependent 1E [Source:HGNC Symbol;Acc:HGNC:19322] |
| **ENSG00000233521** | LINC01638 | 105372978 | 15 | 10.90758268 | -1.281684763 | 0.449492042 | -2.851407019 | 0.004352621 | long intergenic non-protein coding RNA 1638 [Source:HGNC Symbol;Acc:HGNC:52425] |
| **ENSG00000144810** | COL8A1 | 1295 | 14 | 54.90580421 | -0.588464392 | 0.202784134 | -2.90192522 | 0.003708771 | collagen type VIII alpha 1 chain [Source:HGNC Symbol;Acc:HGNC:2215] |
| **ENSG00000225975** | LINC01534 | NA | 13 | 39.14756271 | -0.77519099 | 0.235222012 | -3.29557164 | 0.000982217 | long intergenic non-protein coding RNA 1534 [Source:HGNC Symbol;Acc:HGNC:51281] |
| **ENSG00000227234** | SPANXB1 | 728695 | 12 | 77.23024801 | -0.776982145 | 0.252120292 | -3.081791396 | 0.00205759 | SPANX family member B1 [Source:HGNC Symbol;Acc:HGNC:14329] |
| **ENSG00000159307** | SCUBE1 | 80274 | 11 | 57.73300288 | -0.681191835 | 0.333467529 | -2.04275312 | 0.041076883 | signal peptide, CUB domain and EGF like domain containing 1 [Source:HGNC Symbol;Acc:HGNC:13441] |
| **ENSG00000156453** | PCDH1 | 5097 | 10 | 48.93153738 | -0.734529771 | 0.272001756 | -2.700459671 | 0.006924373 | protocadherin 1 [Source:HGNC Symbol;Acc:HGNC:8655] |
| **ENSG00000170214** | ADRA1B | 147 | 9 | 60.42023195 | -1.108083814 | 0.232184408 | -4.772429917 | 1.82E-06 | adrenoceptor alpha 1B [Source:HGNC Symbol;Acc:HGNC:278] |
| **ENSG00000175040** | CHST2 | 9435 | 8 | 106.3545515 | -0.695439738 | 0.191770616 | -3.626414467 | 0.000287384 | carbohydrate sulfotransferase 2 [Source:HGNC Symbol;Acc:HGNC:1970] |
| **ENSG00000118432** | CNR1 | 1268 | 7 | 5.294011445 | -1.445792029 | 0.589069032 | -2.454367741 | 0.014113257 | cannabinoid receptor 1 [Source:HGNC Symbol;Acc:HGNC:2159] |
| **ENSG00000005249** | PRKAR2B | 5577 | 6 | 37.71235692 | -0.728825768 | 0.285196972 | -2.555517203 | 0.010603012 | protein kinase cAMP-dependent type II regulatory subunit beta [Source:HGNC Symbol;Acc:HGNC:9392] |
| **ENSG00000245146** | MALINC1 | NA | 5 | 13.08402792 | -1.184357832 | 0.387600141 | -3.055617651 | 0.002245975 | mitosis associated long intergenic non-coding RNA 1 [Source:HGNC Symbol;Acc:HGNC:49009] |
| **ENSG00000007372** | PAX6 | 5080 | 4 | 30.35991131 | -0.653146491 | 0.241569151 | -2.703766141 | 0.006855852 | paired box 6 [Source:HGNC Symbol;Acc:HGNC:8620] |
| **ENSG00000162738** | VANGL2 | 57216 | 3 | 38.98412356 | -1.079815539 | 0.272328243 | -3.965125053 | 7.34E-05 | VANGL planar cell polarity protein 2 [Source:HGNC Symbol;Acc:HGNC:15511] |
| **ENSG00000144278** | GALNT13 | 114805 | 2 | 33.16780325 | -0.662831463 | 0.270645742 | -2.449074051 | 0.0143224 | polypeptide N-acetylgalactosaminyltransferase 13 [Source:HGNC Symbol;Acc:HGNC:23242] |
| **ENSG00000157168** | NRG1 | 3084 | 1 | 9.912399205 | -0.983091402 | 0.46764951 | -2.102197014 | 0.035536021 | neuregulin 1 [Source:HGNC Symbol;Acc:HGNC:7997] |
