## Supplemental Figure 2B for "Adenomatous Polyposis Coli Loss Controls Cell Cycle Regulators and Response to Paclitaxel"

Supplemental Figure 2B. Up-regulated in the APC shRNA1 cells

| ensembl_gene_id | hgnc | entrezgene_id | ids | baseMean | log2FoldChange | lfcSE | stat | pvalue | description |
|---|---|---|---|---|---|---|---|---|---|
| ENSG00000125637 | PSD4 | 23550 | 190 | 45.62179794 | 1.660192691 | 0.303440068 | 5.47123753 | 4.47E-08 | pleckstrin and Sec7 domain containing 4 [Source:HGNC Symbol;Acc:HGNC:19096] |
| ENSG00000167191 | GPRC5B | 51704 | 189 | 167.7575926 | 0.652971279 | 0.176337924 | 3.702954324 | 0.000213103 | G protein-coupled receptor class C group 5 member B [Source:HGNC Symbol;Acc:HGNC:13308] |
| ENSG00000107485 | GATA3 | 2625 | 188 | 11.8358029 | 2.908320678 | 0.656747381 | 4.428370424 | 9.49E-06 | GATA binding protein 3 [Source:HGNC Symbol;Acc:HGNC:4172] |
| ENSG00000135924 | DNAJB2 | 3300 | 187 | 1303.22337 | 1.032260733 | 0.153705497 | 6.715834857 | 1.87E-11 | DnaJ heat shock protein family (Hsp40) member B2 [Source:HGNC Symbol;Acc:HGNC:5228] |
| ENSG00000136717 | BIN1 | 274 | 186 | 1299.310413 | 1.059609053 | 0.162447847 | 6.522764506 | 6.9E-11 | bridging integrator 1 [Source:HGNC Symbol;Acc:HGNC:1052] |
| ENSG00000160888 | IER2 | 9592 | 185 | 910.6429647 | 0.961993417 | 0.137220009 | 7.010591421 | 2.37E-12 | immediate early response 2 [Source:HGNC Symbol;Acc:HGNC:28871] |
| ENSG00000197111 | PCBP2 | 5094 | 184 | 1777.330814 | 0.866911125 | 0.13415194 | 6.462158694 | 1.03E-10 | poly(rC) binding protein 2 [Source:HGNC Symbol;Acc:HGNC:8648] |
| ENSG00000182103 | FAM181B | 220382 | 182 | 77.42158705 | 1.600935904 | 0.243823825 | 6.565953531 | 5.17E-11 | family with sequence similarity 181 member B [Source:HGNC Symbol;Acc:HGNC:28512] |
| ENSG00000133265 | HSPBP1 | 23640 | 181 | 4266.375207 | 1.607906019 | 0.240787799 | 6.677688937 | 2.43E-11 | HSPA (Hsp70) binding protein 1 [Source:HGNC Symbol;Acc:HGNC:24989] |
| ENSG00000137331 | IER3 | 8870 | 180 | 1456.42362 | 1.359737234 | 0.193857277 | 7.01411501 | 2.31E-12 | immediate early response 3 [Source:HGNC Symbol;Acc:HGNC:5392] |
| ENSG00000011105 | TSPAN9 | 10867 | 179 | 1856.714042 | 0.999488653 | 0.163670815 | 6.106700544 | 1.02E-09 | tetraspanin 9 [Source:HGNC Symbol;Acc:HGNC:21640] |
| ENSG00000156381 | ANKRD9 | 122416 | 178 | 797.6217251 | 1.183669393 | 0.179527272 | 6.593256735 | 4.3E-11 | ankyrin repeat domain 9 [Source:HGNC Symbol;Acc:HGNC:20096] |
| ENSG00000185838 | GNB1L | 54584 | 177 | 391.030043 | 1.514355374 | 0.21705679 | 6.976770331 | 3.02E-12 | G protein subunit beta 1 like [Source:HGNC Symbol;Acc:HGNC:4397] |
| ENSG00000161249 | DMKN | 93099 | 176 | 145.4444976 | 2.104584578 | 0.207812726 | 10.12731329 | 4.18E-24 | dermokine [Source:HGNC Symbol;Acc:HGNC:25063] |
| ENSG00000111057 | KRT18 | 3875 | 175 | 5691.2309 | 1.50734628 | 0.21142244 | 7.129547272 | 1.01E-12 | keratin 18 [Source:HGNC Symbol;Acc:HGNC:6430] |
| ENSG00000179943 | FIZ1 | 84922 | 174 | 512.6228392 | 1.096999165 | 0.167074301 | 6.565935978 | 5.17E-11 | FLT3 interacting zinc finger 1 [Source:HGNC Symbol;Acc:HGNC:25917] |
| ENSG00000130560 | UBAC1 | 10422 | 173 | 1089.035077 | 0.935638128 | 0.173082154 | 5.405745809 | 6.45E-08 | UBA domain containing 1 [Source:HGNC Symbol;Acc:HGNC:30221] |
| ENSG00000171621 | SPSB1 | 80176 | 172 | 178.8363239 | 0.994261804 | 0.185236302 | 5.367532127 | 7.98E-08 | splA/ryanodine receptor domain and SOCS box containing 1 [Source:HGNC Symbol;Acc:HGNC:30628] |
| ENSG00000143321 | HDGF | 3068 | 171 | 6999.655947 | 1.033125855 | 0.191034668 | 5.408054293 | 6.37E-08 | heparin binding growth factor [Source:HGNC Symbol;Acc:HGNC:4856] |
| ENSG00000107438 | PDLIM1 | 9124 | 170 | 865.1782077 | 0.88881172 | 0.162496345 | 5.469733625 | 4.51E-08 | PDZ and LIM domain 1 [Source:HGNC Symbol;Acc:HGNC:2067] |
| ENSG00000162783 | IER5 | 51278 | 169 | 481.2622447 | 0.928163797 | 0.131730015 | 7.045955303 | 1.84E-12 | immediate early response 5 [Source:HGNC Symbol;Acc:HGNC:5393] |
| ENSG00000163453 | IGFBP7 | 3490 | 168 | 1050.9188 | 1.255526753 | 0.194878224 | 6.442622101 | 1.17E-10 | insulin like growth factor binding protein 7 [Source:HGNC Symbol;Acc:HGNC:5476] |
| ENSG00000140406 | TLNRD1 | 59274 | 167 | 336.6942218 | 0.858728046 | 0.180475572 | 4.75814004 | 1.95E-06 | talin rod domain containing 1 [Source:HGNC Symbol;Acc:HGNC:13519] |
| ENSG00000159403 | C1R | 715 | 166 | 296.1074188 | 0.755463595 | 0.139619535 | 5.410873151 | 6.27E-08 | complement C1r [Source:HGNC Symbol;Acc:HGNC:1246] |
| ENSG00000149532 | CPSF7 | 79869 | 165 | 1176.154659 | 0.600368796 | 0.12727984 | 4.716919805 | 2.39E-06 | cleavage and polyadenylation specific factor 7 [Source:HGNC Symbol;Acc:HGNC:30098] |
| ENSG00000167566 | NCKAP5L | 57701 | 164 | 785.5398031 | 1.099964327 | 0.201031824 | 5.471593027 | 4.46E-08 | NCK associated protein 5 like [Source:HGNC Symbol;Acc:HGNC:29321] |
| ENSG00000180758 | GPR157 | 80045 | 163 | 159.4055622 | 1.434396438 | 0.190778902 | 7.518632423 | 5.54E-14 | G protein-coupled receptor 157 [Source:HGNC Symbol;Acc:HGNC:23687] |
| ENSG00000147799 | ARHGAP39 | 80728 | 162 | 599.6921252 | 0.963451043 | 0.133862807 | 7.197301959 | 6.14E-13 | Rho GTPase activating protein 39 [Source:HGNC Symbol;Acc:HGNC:29351] |
| ENSG00000204388 | HSPA1B | 3304 | 161 | 741.2424179 | 0.996211592 | 0.13624079 | 7.312138966 | 2.63E-13 | heat shock protein family A (Hsp70) member 1B [Source:HGNC Symbol;Acc:HGNC:5233] |
| ENSG00000102572 | STK24 | 8428 | 160 | 1064.780836 | 1.047225132 | 0.132584101 | 7.89857248 | 2.82E-15 | serine/threonine kinase 24 [Source:HGNC Symbol;Acc:HGNC:11403] |
| ENSG00000170421 | KRT8 | 3856 | 159 | 11184.29519 | 2.057155848 | 0.216173657 | 9.516218916 | 1.8E-21 | keratin 8 [Source:HGNC Symbol;Acc:HGNC:6446] |
| ENSG00000088247 | KHSRP | 8570 | 158 | 6613.117588 | 1.010172278 | 0.141159739 | 7.156235112 | 8.29E-13 | KH-type splicing regulatory protein [Source:HGNC Symbol;Acc:HGNC:6316] |
| ENSG00000063244 | U2AF2 | 11338 | 157 | 2411.822057 | 1.089985736 | 0.158479517 | 6.877770442 | 6.08E-12 | U2 small nuclear RNA auxiliary factor 2 [Source:HGNC Symbol;Acc:HGNC:23156] |
| ENSG00000143878 | RHOB | 388 | 156 | 1803.561265 | 1.353360115 | 0.161035654 | 8.404102315 | 4.31E-17 | ras homolog family member B [Source:HGNC Symbol;Acc:HGNC:668] |
| ENSG00000127663 | KDM4B | 23030 | 155 | 1254.544064 | 1.139031722 | 0.151709371 | 7.507985253 | 6E-14 | lysine demethylase 4B [Source:HGNC Symbol;Acc:HGNC:29136] |
| ENSG00000005007 | UPF1 | 5976 | 154 | 2628.547066 | 1.106049532 | 0.129435158 | 8.545201718 | 1.28E-17 | UPF1 RNA helicase and ATPase [Source:HGNC Symbol;Acc:HGNC:9962] |
| ENSG00000146830 | GIGYF1 | 64599 | 153 | 602.5988307 | 0.941128597 | 0.172957495 | 5.44138662 | 5.29E-08 | GRB10 interacting GYF protein 1 [Source:HGNC Symbol;Acc:HGNC:9126] |
| ENSG00000176046 | NUPR1 | 26471 | 152 | 38.40806003 | 2.780059461 | 0.339889662 | 8.179299839 | 2.85E-16 | nuclear protein 1, transcriptional regulator [Source:HGNC Symbol;Acc:HGNC:29990] |
| ENSG00000080573 | COL5A3 | 50509 | 151 | 2864.394665 | 1.276319312 | 0.196288331 | 6.502267883 | 7.91E-11 | collagen type V alpha 3 chain [Source:HGNC Symbol;Acc:HGNC:14864] |
| ENSG00000167491 | GATAD2A | 54815 | 150 | 1744.832396 | 0.739182419 | 0.11351198 | 6.511933072 | 7.42E-11 | GATA zinc finger domain containing 2A [Source:HGNC Symbol;Acc:HGNC:29989] |
| ENSG00000149257 | SERPINH1 | 871 | 149 | 687.1096701 | 1.549234094 | 0.153722393 | 10.07812891 | 6.9E-24 | serpin family H member 1 [Source:HGNC Symbol;Acc:HGNC:1546] |
| ENSG00000165801 | ARHGEF40 | 55701 | 148 | 1518.793642 | 0.966872332 | 0.16867194 | 5.732265449 | 9.91E-09 | Rho guanine nucleotide exchange factor 40 [Source:HGNC Symbol;Acc:HGNC:25516] |
| ENSG00000130382 | MLLT1 | 4298 | 147 | 1991.273017 | 0.94531464 | 0.141414864 | 6.684690811 | 2.31E-11 | MLLT1 super elongation complex subunit [Source:HGNC Symbol;Acc:HGNC:7134] |
| ENSG00000197622 | CDC42SE1 | 56882 | 146 | 438.2227431 | 0.741461416 | 0.12448065 | 5.956439144 | 2.58E-09 | CDC42 small effector 1 [Source:HGNC Symbol;Acc:HGNC:17719] |
| ENSG00000105173 | CCNE1 | 898 | 145 | 1767.45658 | 0.933998372 | 0.118399365 | 7.888542072 | 3.06E-15 | cyclin E1 [Source:HGNC Symbol;Acc:HGNC:1589] |
| ENSG00000137076 | TLN1 | 7094 | 144 | 7247.246479 | 1.019505907 | 0.181347538 | 5.621834832 | 1.89E-08 | talin 1 [Source:HGNC Symbol;Acc:HGNC:11845] |
| ENSG00000022556 | NLRP2 | 55655 | 143 | 1668.020727 | 1.115202414 | 0.137253706 | 8.125116952 | 4.47E-16 | NLR family pyrin domain containing 2 [Source:HGNC Symbol;Acc:HGNC:22948] |
| ENSG00000163874 | ZC3H12A | 80149 | 142 | 116.595613 | 1.273498784 | 0.186660434 | 6.822542698 | 8.94E-12 | zinc finger CCCH-type containing 12A [Source:HGNC Symbol;Acc:HGNC:26259] |
| ENSG00000181090 | EHMT1 | 79813 | 141 | 515.3936471 | 0.901689932 | 0.123234551 | 7.316859819 | 2.54E-13 | euchromatic histone lysine methyltransferase 1 [Source:HGNC Symbol;Acc:HGNC:24650] |
| ENSG00000068654 | POLR1A | 25885 | 140 | 1649.936837 | 0.818518279 | 0.125843102 | 6.504276105 | 7.81E-11 | RNA polymerase I subunit A [Source:HGNC Symbol;Acc:HGNC:17264] |
| ENSG00000158195 | WASF2 | 10163 | 139 | 2319.546168 | 0.804294252 | 0.147685148 | 5.446006338 | 5.15E-08 | WASP family member 2 [Source:HGNC Symbol;Acc:HGNC:12733] |
| ENSG00000130402 | ACTN4 | 81 | 138 | 15874.75795 | 1.082718285 | 0.14809047 | 7.311194887 | 2.65E-13 | actinin alpha 4 [Source:HGNC Symbol;Acc:HGNC:166] |
| ENSG00000095383 | TBC1D2 | 55357 | 137 | 543.688807 | 1.082627753 | 0.160022705 | 6.765463376 | 1.33E-11 | TBC1 domain family member 2 [Source:HGNC Symbol;Acc:HGNC:18026] |
| ENSG00000204389 | HSPA1A | 3303 | 136 | 648.155884 | 1.051281021 | 0.155124304 | 6.77702328 | 1.23E-11 | heat shock protein family A (Hsp70) member 1A [Source:HGNC Symbol;Acc:HGNC:5232] |
| ENSG00000127527 | EPS15L1 | 58513 | 135 | 699.5602713 | 0.901388879 | 0.131439723 | 6.857811795 | 6.99E-12 | epidermal growth factor receptor pathway substrate 15 like 1 [Source:HGNC Symbol;Acc:HGNC:24634] |
| ENSG00000167658 | EEF2 | 1938 | 134 | 31866.47468 | 1.334498278 | 0.206356331 | 6.466960664 | 1E-10 | eukaryotic translation elongation factor 2 [Source:HGNC Symbol;Acc:HGNC:3214] |
| ENSG00000149115 | TNKS1BP1 | 85456 | 133 | 1531.399495 | 0.954432387 | 0.155365363 | 6.143147797 | 8.09E-10 | tankyrase 1 binding protein 1 [Source:HGNC Symbol;Acc:HGNC:19081] |
| ENSG00000168487 | BMP1 | 649 | 132 | 1311.931608 | 0.929367806 | 0.157083445 | 5.916395616 | 3.29E-09 | bone morphogenetic protein 1 [Source:HGNC Symbol;Acc:HGNC:1067] |
| ENSG00000136699 | SMPD4 | 55627 | 131 | 1370.301235 | 0.877320688 | 0.132146123 | 6.6390195 | 3.16E-11 | sphingomyelin phosphodiesterase 4 [Source:HGNC Symbol;Acc:HGNC:32949] |
| ENSG00000011243 | AKAP8L | 26993 | 130 | 539.665454 | 0.931400348 | 0.113649391 | 8.195383548 | 2.5E-16 | A-kinase anchoring protein 8 like [Source:HGNC Symbol;Acc:HGNC:29857] |
| ENSG00000125733 | TRIP10 | 9322 | 129 | 1064.294161 | 1.171241461 | 0.150098524 | 7.803151064 | 6.04E-15 | thyroid hormone receptor interactor 10 [Source:HGNC Symbol;Acc:HGNC:12304] |
| ENSG00000125731 | SH2D3A | 10045 | 128 | 318.2263808 | 1.29715983 | 0.209635139 | 6.187702294 | 6.1E-10 | SH2 domain containing 3A [Source:HGNC Symbol;Acc:HGNC:16885] |
| ENSG00000070047 | PHRF1 | 57661 | 127 | 1089.174035 | 0.7878242 | 0.143656781 | 5.484072483 | 4.16E-08 | PHD and ring finger domains 1 [Source:HGNC Symbol;Acc:HGNC:24351] |
| ENSG00000160877 | NACC1 | 112939 | 126 | 3220.685472 | 0.97775185 | 0.211869911 | 4.614868842 | 3.93E-06 | nucleus accumbens associated 1 [Source:HGNC Symbol;Acc:HGNC:20967] |
| ENSG00000072609 | CHFR | 55743 | 125 | 707.7149467 | 0.8054214 | 0.164679028 | 4.890855923 | 1E-06 | checkpoint with forkhead and ring finger domains [Source:HGNC Symbol;Acc:HGNC:20455] |
| ENSG00000185252 | ZNF74 | 7625 | 124 | 358.1845307 | 1.004385131 | 0.15252455 | 6.585071919 | 4.55E-11 | zinc finger protein 74 [Source:HGNC Symbol;Acc:HGNC:13144] |
| ENSG00000162413 | KLHL21 | 9903 | 123 | 996.9344953 | 0.997100991 | 0.184918569 | 5.392108508 | 6.96E-08 | kelch like family member 21 [Source:HGNC Symbol;Acc:HGNC:29041] |
| ENSG00000092203 | TOX4 | 9878 | 122 | 1281.854095 | 0.853191109 | 0.139216999 | 6.128498064 | 8.87E-10 | TOX high mobility group box family member 4 [Source:HGNC Symbol;Acc:HGNC:20161] |
| ENSG00000197951 | ZNF71 | 58491 | 121 | 914.5294157 | 0.756456742 | 0.158490978 | 4.772869413 | 1.82E-06 | zinc finger protein 71 [Source:HGNC Symbol;Acc:HGNC:13141] |
| ENSG00000107731 | UNC5B | 219699 | 120 | 80.36449343 | 1.306155109 | 0.238547343 | 5.475454447 | 4.36E-08 | unc-5 netrin receptor B [Source:HGNC Symbol;Acc:HGNC:12568] |
| ENSG00000135912 | TTLL4 | 9654 | 119 | 457.6656224 | 0.787611499 | 0.114953261 | 6.851580304 | 7.3E-12 | tubulin tyrosine ligase like 4 [Source:HGNC Symbol;Acc:HGNC:28976] |
| ENSG00000118515 | SGK1 | 6446 | 118 | 330.7357534 | 1.563926777 | 0.13846923 | 11.29439932 | 1.4E-29 | serum/glucocorticoid regulated kinase 1 [Source:HGNC Symbol;Acc:HGNC:10810] |
| ENSG00000115844 | DLX2 | 1746 | 117 | 264.1369877 | 1.201105157 | 0.120053186 | 10.00477536 | 1.45E-23 | distal-less homeobox 2 [Source:HGNC Symbol;Acc:HGNC:2915] |
| ENSG00000124225 | PMEPA1 | 56937 | 116 | 1145.495416 | 1.074854357 | 0.101826117 | 10.5557826 | 4.78E-26 | prostate transmembrane protein, androgen induced 1 [Source:HGNC Symbol;Acc:HGNC:14107] |

| | | | | | | | | | |
|---|---|---|---|---|---|---|---|---|---|
| ENSG00000104892 | KLC3 | 147700 | 115 | 144.7803119 | 2.171451177 | 0.297258609 | 7.304922753 | 2.77E-13 | kinesin light chain 3 [Source:HGNC Symbol;Acc:HGNC:20717] |
| ENSG00000152137 | HSPB8 | 26353 | 114 | 539.2141766 | 1.59380694 | 0.145399198 | 10.96159374 | 5.85E-28 | heat shock protein family B (small) member 8 [Source:HGNC Symbol;Acc:HGNC:30171] |
| ENSG00000182578 | CSF1R | 1436 | 113 | 150.0862315 | 2.293112081 | 0.214798273 | 10.67565417 | 1.32E-26 | colony stimulating factor 1 receptor [Source:HGNC Symbol;Acc:HGNC:2433] |
| ENSG00000179528 | LBX2 | 85474 | 112 | 63.87962149 | 1.346400005 | 0.204460815 | 6.585124905 | 4.55E-11 | ladybird homeobox 2 [Source:HGNC Symbol;Acc:HGNC:15525] |
| ENSG00000129354 | AP1M2 | 10053 | 111 | 536.8694424 | 3.403717457 | 0.232371252 | 14.64775622 | 1.39E-48 | adaptor related protein complex 1 subunit mu 2 [Source:HGNC Symbol;Acc:HGNC:558] |
| ENSG00000149591 | TAGLN | 6876 | 110 | 115.4047899 | 2.28427012 | 0.251174136 | 9.094368377 | 9.51E-20 | transgelin [Source:HGNC Symbol;Acc:HGNC:11553] |
| ENSG00000232677 | LINC00665 | NA | 109 | 747.9304703 | 0.917013183 | 0.142344248 | 6.44222155 | 1.18E-10 | long intergenic non-protein coding RNA 665 [Source:HGNC Symbol;Acc:HGNC:44323] |
| ENSG00000185112 | FAM43A | 131583 | 108 | 310.8169162 | 1.189292041 | 0.164300505 | 7.238517242 | 4.54E-13 | family with sequence similarity 43 member A [Source:HGNC Symbol;Acc:HGNC:26888] |
| ENSG00000167470 | MIDN | 90007 | 107 | 591.7008719 | 1.03868903 | 0.148315698 | 7.003230562 | 2.5E-12 | midnolin [Source:HGNC Symbol;Acc:HGNC:16298] |
| ENSG00000171853 | TRAPPC12 | 51112 | 106 | 782.9192803 | 0.691344937 | 0.10501994 | 6.582987366 | 4.61E-11 | trafficking protein particle complex 12 [Source:HGNC Symbol;Acc:HGNC:24284] |
| ENSG00000112787 | FBRSL1 | 57666 | 105 | 744.1731209 | 1.041128208 | 0.164876003 | 6.314613339 | 2.71E-10 | fibrosin like 1 [Source:HGNC Symbol;Acc:HGNC:29308] |
| ENSG00000134030 | CTIF | 9811 | 104 | 360.9993854 | 1.157059414 | 0.178725328 | 6.473953231 | 9.55E-11 | cap binding complex dependent translation initiation factor [Source:HGNC Symbol;Acc:HGNC:23925] |
| ENSG00000010278 | CD9 | 928 | 103 | 2414.290141 | 0.982103466 | 0.127780312 | 7.68587469 | 1.52E-14 | CD9 molecule [Source:HGNC Symbol;Acc:HGNC:1709] |
| ENSG00000143772 | ITPKB | 3707 | 102 | 551.5100192 | 1.224789968 | 0.169351667 | 7.232228576 | 4.75E-13 | inositol-trisphosphate 3-kinase B [Source:HGNC Symbol;Acc:HGNC:6179] |
| ENSG00000100814 | CCNB1IP1 | 57820 | 101 | 452.8707302 | 1.079276361 | 0.138249354 | 7.806737107 | 5.87E-15 | cyclin B1 interacting protein 1 [Source:HGNC Symbol;Acc:HGNC:19437] |
| ENSG00000138092 | CENPO | 79172 | 100 | 407.0570619 | 0.834453331 | 0.127115437 | 6.564531817 | 5.22E-11 | centromere protein O [Source:HGNC Symbol;Acc:HGNC:28152] |
| ENSG00000176871 | WSB2 | 55884 | 99 | 845.5937797 | 0.863633617 | 0.120679588 | 7.156418347 | 8.28E-13 | WD repeat and SOCS box containing 2 [Source:HGNC Symbol;Acc:HGNC:19222] |
| ENSG00000110042 | DTX4 | 23220 | 98 | 254.3651069 | 1.20896722 | 0.152613042 | 7.921781804 | 2.34E-15 | deltex E3 ubiquitin ligase 4 [Source:HGNC Symbol;Acc:HGNC:29151] |
| ENSG00000166908 | PIP4K2C | 79837 | 97 | 604.9381354 | 0.84945359 | 0.118982834 | 7.139295314 | 9.38E-13 | phosphatidylinositol-5-phosphate 4-kinase type 2 gamma [Source:HGNC Symbol;Acc:HGNC:23786] |
| ENSG00000116017 | ARID3A | 1820 | 96 | 194.4276165 | 0.692102123 | 0.12919195 | 5.357161365 | 8.45E-08 | AT-rich interaction domain 3A [Source:HGNC Symbol;Acc:HGNC:3031] |
| ENSG00000143614 | GATAD2B | 57459 | 95 | 306.3068155 | 1.077411234 | 0.146421742 | 7.35827357 | 1.86E-13 | GATA zinc finger domain containing 2B [Source:HGNC Symbol;Acc:HGNC:30778] |
| ENSG00000172061 | LRRC15 | 131578 | 93 | 22.24335376 | 1.262599204 | 0.410947172 | 3.072412443 | 0.002123361 | leucine rich repeat containing 15 [Source:HGNC Symbol;Acc:HGNC:20818] |
| ENSG00000143147 | GPR161 | 23432 | 92 | 509.1292255 | 0.693606906 | 0.106043302 | 6.540789391 | 6.12E-11 | G protein-coupled receptor 161 [Source:HGNC Symbol;Acc:HGNC:23694] |
| ENSG00000074047 | GLI2 | 2736 | 91 | 260.275435 | 1.057268164 | 0.135561967 | 7.799150364 | 6.23E-15 | GLI family zinc finger 2 [Source:HGNC Symbol;Acc:HGNC:4318] |
| ENSG00000135525 | MAP7 | 9053 | 90 | 64.93662682 | 1.241539863 | 0.243111705 | 5.106869966 | 3.28E-07 | microtubule associated protein 7 [Source:HGNC Symbol;Acc:HGNC:6869] |
| ENSG00000137460 | FHDC1 | 85462 | 89 | 148.2468776 | 0.850269764 | 0.138376916 | 6.144592522 | 8.02E-10 | FH2 domain containing 1 [Source:HGNC Symbol;Acc:HGNC:29363] |
| ENSG00000160439 | RDH13 | 112724 | 88 | 78.81327864 | 1.491986578 | 0.229832268 | 6.491632304 | 8.49E-11 | retinol dehydrogenase 13 [Source:HGNC Symbol;Acc:HGNC:19978] |
| ENSG00000065970 | FOXJ2 | 55810 | 87 | 433.9356574 | 0.953713376 | 0.11020858 | 8.6537126 | 4.99E-18 | forkhead box J2 [Source:HGNC Symbol;Acc:HGNC:24818] |
| ENSG00000115718 | PROC | 5624 | 86 | 85.41013936 | 1.751627585 | 0.211715288 | 8.273505428 | 1.3E-16 | protein C, inactivator of coagulation factors Va and VIIIa [Source:HGNC Symbol;Acc:HGNC:9451] |
| ENSG00000084092 | NOA1 | 84273 | 85 | 208.3261986 | 0.749601456 | 0.110880037 | 6.760472629 | 1.38E-11 | nitric oxide associated 1 [Source:HGNC Symbol;Acc:HGNC:28473] |
| ENSG00000125730 | C3 | 718 | 84 | 138.4209353 | 2.720455203 | 0.205586638 | 13.23264597 | 5.69E-40 | complement C3 [Source:HGNC Symbol;Acc:HGNC:1318] |
| ENSG00000130803 | ZNF317 | 57693 | 83 | 723.5033937 | 0.78072636 | 0.142866725 | 5.464717955 | 4.64E-08 | zinc finger protein 317 [Source:HGNC Symbol;Acc:HGNC:13507] |
| ENSG00000099194 | SCD | 6319 | 82 | 1062.046411 | 1.145640948 | 0.12641425 | 9.062593384 | 1.27E-19 | stearoyl-CoA desaturase [Source:HGNC Symbol;Acc:HGNC:10571] |
| ENSG00000061936 | SFSWAP | 6433 | 81 | 483.4566095 | 0.692298114 | 0.100595202 | 6.882019232 | 5.9E-12 | splicing factor SWAP [Source:HGNC Symbol;Acc:HGNC:10790] |
| ENSG00000170390 | DCLK2 | 166614 | 80 | 319.2411877 | 1.098210437 | 0.115031341 | 9.547054098 | 1.33E-21 | doublecortin like kinase 2 [Source:HGNC Symbol;Acc:HGNC:19002] |
| ENSG00000064195 | DLX3 | 1747 | 79 | 59.42044107 | 4.988673607 | 0.38213048 | 13.05489581 | 5.96E-39 | distal-less homeobox 3 [Source:HGNC Symbol;Acc:HGNC:2916] |
| ENSG00000185585 | OLFML2A | 169611 | 78 | 504.344028 | 1.204466845 | 0.155524375 | 7.744553522 | 9.59E-15 | olfactomedin like 2A [Source:HGNC Symbol;Acc:HGNC:27270] |
| ENSG00000213626 | LBH | 81606 | 77 | 393.9340595 | 1.933618283 | 0.158307491 | 12.21431958 | 2.61E-34 | LBH regulator of WNT signaling pathway [Source:HGNC Symbol;Acc:HGNC:29532] |
| ENSG00000168542 | COL3A1 | 1281 | 76 | 4242.726117 | 1.217824864 | 0.13612776 | 8.94619044 | 3.68E-19 | collagen type III alpha 1 chain [Source:HGNC Symbol;Acc:HGNC:2201] |
| ENSG00000099331 | MYO9B | 4650 | 75 | 1943.136863 | 1.007672637 | 0.150038956 | 6.716073361 | 1.87E-11 | myosin IXB [Source:HGNC Symbol;Acc:HGNC:7609] |
| ENSG00000075426 | FOSL2 | 2355 | 74 | 1677.466794 | 0.794340015 | 0.114332568 | 6.947626843 | 3.71E-12 | FOS like 2, AP-1 transcription factor subunit [Source:HGNC Symbol;Acc:HGNC:3798] |
| ENSG00000108821 | COL1A1 | 1277 | 73 | 13428.53979 | 2.278951309 | 0.151990583 | 14.99402963 | 8.03E-51 | collagen type I alpha 1 chain [Source:HGNC Symbol;Acc:HGNC:2197] |
| ENSG00000132359 | RAP1GAP2 | 23108 | 72 | 81.4164133 | 1.410042851 | 0.185357024 | 7.607172485 | 2.8E-14 | RAP1 GTPase activating protein 2 [Source:HGNC Symbol;Acc:HGNC:29176] |
| ENSG00000177084 | POLE | 5426 | 71 | 1541.387807 | 0.741268572 | 0.122353763 | 6.058404372 | 1.37E-09 | DNA polymerase epsilon, catalytic subunit [Source:HGNC Symbol;Acc:HGNC:9177] |
| ENSG00000114270 | COL7A1 | 1294 | 70 | 299.3661705 | 1.480704195 | 0.171031315 | 8.657503419 | 4.82E-18 | collagen type VII alpha 1 chain [Source:HGNC Symbol;Acc:HGNC:2214] |
| ENSG00000173801 | JUP | 3728 | 69 | 1187.589784 | 1.721318691 | 0.204134061 | 8.432295347 | 3.39E-17 | junction plakoglobin [Source:HGNC Symbol;Acc:HGNC:6207] |
| ENSG00000155275 | TRMT44 | 152992 | 68 | 136.0203497 | 0.802420026 | 0.147850411 | 5.427242446 | 5.72E-08 | tRNA methyltransferase 44 homolog [Source:HGNC Symbol;Acc:HGNC:26653] |
| ENSG00000130635 | COL5A1 | 1289 | 67 | 6681.072462 | 0.891815812 | 0.163942081 | 5.439822441 | 5.33E-08 | collagen type V alpha 1 chain [Source:HGNC Symbol;Acc:HGNC:2209] |
| ENSG00000089154 | GCN1 | 10985 | 66 | 3660.723974 | 0.961292086 | 0.166200578 | 5.783927454 | 7.3E-09 | GCN1 activator of EIF2AK4 [Source:HGNC Symbol;Acc:HGNC:4199] |
| ENSG00000272333 | KMT2B | 9757 | 65 | 2019.499207 | 0.965050699 | 0.170905841 | 5.646680614 | 1.64E-08 | lysine methyltransferase 2B [Source:HGNC Symbol;Acc:HGNC:15840] |
| ENSG00000178209 | PLEC | 5339 | 64 | 17189.12856 | 1.216142565 | 0.178793634 | 6.801934388 | 1.03E-11 | plectin [Source:HGNC Symbol;Acc:HGNC:9069] |
| ENSG00000196498 | NCOR2 | 9612 | 63 | 2989.320989 | 1.22428167 | 0.141464278 | 8.654352114 | 4.96E-18 | nuclear receptor corepressor 2 [Source:HGNC Symbol;Acc:HGNC:7673] |
| ENSG00000143367 | TUFT1 | 7286 | 62 | 440.9239551 | 1.109804942 | 0.139412166 | 7.960603255 | 1.71E-15 | tuftelin 1 [Source:HGNC Symbol;Acc:HGNC:12422] |
| ENSG00000100181 | TPTEP1 | NA | 61 | 269.1399771 | 1.246449786 | 0.1956066 | 6.372227646 | 1.86E-10 | TPTE pseudogene 1 [Source:HGNC Symbol;Acc:HGNC:43648] |
| ENSG00000112139 | MDGA1 | 266727 | 60 | 309.5652374 | 0.788042447 | 0.18067075 | 4.361759976 | 1.29E-05 | MAM domain containing glycosylphosphatidylinositol anchor 1 [Source:HGNC Symbol;Acc:HGNC:19267 |
| ENSG00000197852 | INKA2 | 55924 | 59 | 103.3970802 | 1.436885167 | 0.173154252 | 8.298295622 | 1.06E-16 | inka box actin regulator 2 [Source:HGNC Symbol;Acc:HGNC:28045] |
| ENSG00000108846 | ABCC3 | 8714 | 58 | 229.4728387 | 1.070544785 | 0.144530846 | 7.407033276 | 1.29E-13 | ATP binding cassette subfamily C member 3 [Source:HGNC Symbol;Acc:HGNC:54] |
| ENSG00000261221 | ZNF865 | 100507290 | 57 | 441.9159814 | 0.810628111 | 0.147735136 | 5.487036674 | 4.09E-08 | zinc finger protein 865 [Source:HGNC Symbol;Acc:HGNC:38705] |
| ENSG00000164045 | CDC25A | 993 | 56 | 278.9716048 | 0.882186013 | 0.160804267 | 5.486085852 | 4.11E-08 | cell division cycle 25A [Source:HGNC Symbol;Acc:HGNC:1725] |
| ENSG00000119242 | CCDC92 | 80212 | 55 | 340.8477979 | 0.952396665 | 0.117775774 | 8.086524369 | 6.14E-16 | coiled-coil domain containing 92 [Source:HGNC Symbol;Acc:HGNC:29563] |
| ENSG00000139722 | VPS37B | 79720 | 54 | 151.1529658 | 0.966868276 | 0.220759345 | 4.379738835 | 1.19E-05 | VPS37B subunit of ESCRT-I [Source:HGNC Symbol;Acc:HGNC:25754] |
| ENSG00000197261 | C6orf141 | 135398 | 53 | 29.62628739 | 1.402333461 | 0.319447676 | 4.389869037 | 1.13E-05 | chromosome 6 open reading frame 141 [Source:HGNC Symbol;Acc:HGNC:21351] |
| ENSG00000108819 | PPP1R9B | 84687 | 52 | 370.0997182 | 1.093376704 | 0.12704216 | 8.606408352 | 7.54E-18 | protein phosphatase 1 regulatory subunit 9B [Source:HGNC Symbol;Acc:HGNC:9298] |
| ENSG00000128965 | CHAC1 | 79094 | 51 | 142.2079899 | 1.590197681 | 0.206773824 | 7.690517345 | 1.47E-14 | ChaC glutathione specific gamma-glutamylcyclotransferase 1 [Source:HGNC Symbol;Acc:HGNC:28680] |
| ENSG00000078053 | AMPH | 273 | 50 | 6.69595942 | 2.181220151 | 0.539070486 | 4.046261496 | 5.2E-05 | amphiphysin [Source:HGNC Symbol;Acc:HGNC:471] |
| ENSG00000111087 | GLI1 | 2735 | 49 | 7.078567159 | 1.989019126 | 0.629371331 | 3.160326868 | 0.001575922 | GLI family zinc finger 1 [Source:HGNC Symbol;Acc:HGNC:4317] |
| ENSG00000177663 | IL17RA | 23765 | 48 | 366.6536655 | 0.751653707 | 0.116527133 | 6.450460842 | 1.12E-10 | interleukin 17 receptor A [Source:HGNC Symbol;Acc:HGNC:5985] |
| ENSG00000184545 | DUSP8 | 1850 | 47 | 222.6032711 | 1.413796358 | 0.191044136 | 7.400365111 | 1.36E-13 | dual specificity phosphatase 8 [Source:HGNC Symbol;Acc:HGNC:3074] |
| ENSG00000117318 | ID3 | 3399 | 46 | 2020.029432 | 1.293386902 | 0.207746958 | 6.225780226 | 4.79E-10 | inhibitor of DNA binding 3, HLH protein [Source:HGNC Symbol;Acc:HGNC:5362] |
| ENSG00000129757 | CDKN1C | 1028 | 45 | 234.1320258 | 1.318604603 | 0.209773985 | 6.28583474 | 3.26E-10 | cyclin dependent kinase inhibitor 1C [Source:HGNC Symbol;Acc:HGNC:1786] |
| ENSG00000218891 | ZNF579 | 163033 | 44 | 296.6102983 | 1.063731382 | 0.219839368 | 4.838675581 | 1.31E-06 | zinc finger protein 579 [Source:HGNC Symbol;Acc:HGNC:26646] |
| ENSG00000161956 | SENP3 | 26168 | 43 | 306.344743 | 0.942181319 | 0.206356444 | 4.565795473 | 4.98E-06 | SUMO specific peptidase 3 [Source:HGNC Symbol;Acc:HGNC:17862] |
| ENSG00000154118 | JPH3 | 57338 | 42 | 218.0327488 | 1.238019506 | 0.152967829 | 8.093332514 | 5.81E-16 | junctophilin 3 [Source:HGNC Symbol;Acc:HGNC:14203] |
| ENSG00000184363 | PKP3 | 11187 | 41 | 150.9534706 | 1.293751562 | 0.233832377 | 5.532816199 | 3.15E-08 | plakophilin 3 [Source:HGNC Symbol;Acc:HGNC:9025] |
| ENSG00000104859 | CLASRP | 11129 | 40 | 338.1871868 | 0.819740229 | 0.151158494 | 5.423051042 | 5.86E-08 | CLK4 associating serine/arginine rich protein [Source:HGNC Symbol;Acc:HGNC:17731] |
| ENSG00000133687 | TMTC1 | 83857 | 39 | 40.69140566 | 2.653723656 | 0.28983026 | 9.15613042 | 5.38E-20 | transmembrane O-mannosyltransferase targeting cadherins 1 [Source:HGNC Symbol;Acc:HGNC:24099] |

| | | | | | | | | | |
|---|---|---|---|---|---|---|---|---|---|
| **ENSG00000104369** | JPH1 | 56704 | 38 | 16.53881553 | 3.212844046 | 0.375316396 | 8.560361554 | 1.13E-17 | junctophilin 1 [Source:HGNC Symbol;Acc:HGNC:14201] |
| **ENSG00000135045** | C9orf40 | 55071 | 37 | 213.3245278 | 0.826044072 | 0.130252163 | 6.341883701 | 2.27E-10 | chromosome 9 open reading frame 40 [Source:HGNC Symbol;Acc:HGNC:23433] |
| **ENSG00000127080** | IPPK | 64768 | 36 | 132.4965045 | 0.966092741 | 0.148157037 | 6.520734773 | 7E-11 | inositol-pentakisphosphate 2-kinase [Source:HGNC Symbol;Acc:HGNC:14645] |
| **ENSG00000026559** | KCNG1 | 3755 | 35 | 112.3700685 | 1.282755789 | 0.192102203 | 6.677465259 | 2.43E-11 | potassium voltage-gated channel modifier subfamily G member 1 [Source:HGNC Symbol;Acc:HGNC:624 |
| **ENSG00000170775** | GPR37 | 2861 | 34 | 76.09740216 | 2.47113435 | 0.1970425 | 12.54112359 | 4.45E-36 | G protein-coupled receptor 37 [Source:HGNC Symbol;Acc:HGNC:4494] |
| **ENSG00000125850** | OVOL2 | 58495 | 33 | 9.091588028 | 2.585898117 | 0.575754998 | 4.49131684 | 7.08E-06 | ovo like zinc finger 2 [Source:HGNC Symbol;Acc:HGNC:15804] |
| **ENSG00000004478** | FKBP4 | 2288 | 32 | 103.3631834 | 1.885065625 | 0.250077675 | 7.537920483 | 4.78E-14 | FKBP prolyl isomerase 4 [Source:HGNC Symbol;Acc:HGNC:3720] |
| **ENSG00000163638** | ADAMTS9 | 56999 | 31 | 3.412174648 | 5.233127714 | 1.176507852 | 4.44801767 | 8.67E-06 | ADAM metallopeptidase with thrombospondin type 1 motif 9 [Source:HGNC Symbol;Acc:HGNC:13202] |
| **ENSG00000173334** | TRIB1 | 10221 | 30 | 178.6359857 | 1.045881476 | 0.123061046 | 8.498883362 | 1.91E-17 | tribbles pseudokinase 1 [Source:HGNC Symbol;Acc:HGNC:16891] |
| **ENSG00000141448** | GATA6 | 2627 | 29 | 25.36469883 | 1.764845179 | 0.324726199 | 5.434871541 | 5.48E-08 | GATA binding protein 6 [Source:HGNC Symbol;Acc:HGNC:4174] |
| **ENSG00000179772** | FOXS1 | 2307 | 28 | 3.000706109 | 5.131510673 | 1.372356683 | 3.73919604 | 0.00018461 | forkhead box S1 [Source:HGNC Symbol;Acc:HGNC:3735] |
| **ENSG00000168758** | SEMA4C | 54910 | 27 | 301.4486508 | 0.638261605 | 0.158473935 | 4.027549417 | 5.64E-05 | semaphorin 4C [Source:HGNC Symbol;Acc:HGNC:10731] |
| **ENSG00000167311** | ART5 | 116969 | 26 | 188.0959515 | 0.985842425 | 0.22048902 | 4.471163342 | 7.78E-06 | ADP-ribosyltransferase 5 [Source:HGNC Symbol;Acc:HGNC:24049] |
| **ENSG00000164142** | FAM160A1 | 729830 | 25 | 57.82135282 | 1.49260491 | 0.212905464 | 7.010646346 | 2.37E-12 | family with sequence similarity 160 member A1 [Source:HGNC Symbol;Acc:HGNC:34237] |
| **ENSG00000101384** | JAG1 | 182 | 24 | 458.7934686 | 1.130904856 | 0.138761258 | 8.150004331 | 3.64E-16 | jagged canonical Notch ligand 1 [Source:HGNC Symbol;Acc:HGNC:6188] |
| **ENSG00000101935** | AMMECR1 | 9949 | 23 | 73.17997069 | 1.422922915 | 0.244547175 | 5.818602943 | 5.93E-09 | AMMECR nuclear protein 1 [Source:HGNC Symbol;Acc:HGNC:467] |
| **ENSG00000127481** | UBR4 | 23352 | 22 | 3547.727977 | 0.985001824 | 0.187840628 | 5.243816697 | 1.57E-07 | ubiquitin protein ligase E3 component n-recognin 4 [Source:HGNC Symbol;Acc:HGNC:30313] |
| **ENSG00000166147** | FBN1 | 2200 | 21 | 194.4033605 | 1.346182529 | 0.211769459 | 6.356830379 | 2.06E-10 | fibrillin 1 [Source:HGNC Symbol;Acc:HGNC:3603] |
| **ENSG00000131089** | ARHGEF9 | 23229 | 20 | 54.45171444 | 0.848474027 | 0.198047564 | 4.284193206 | 1.83E-05 | Cdc42 guanine nucleotide exchange factor 9 [Source:HGNC Symbol;Acc:HGNC:14561] |
| **ENSG00000140443** | IGF1R | 3480 | 19 | 167.0156969 | 0.902804039 | 0.170046709 | 5.309153261 | 1.1E-07 | insulin like growth factor 1 receptor [Source:HGNC Symbol;Acc:HGNC:5465] |
| **ENSG00000189221** | MAOA | 4128 | 18 | 7.734354412 | 2.114090704 | 0.511408088 | 4.133862473 | 3.57E-05 | monoamine oxidase A [Source:HGNC Symbol;Acc:HGNC:6833] |
| **ENSG00000198807** | PAX9 | 5083 | 17 | 14.53664829 | 1.770484949 | 0.414445093 | 4.27194091 | 1.94E-05 | paired box 9 [Source:HGNC Symbol;Acc:HGNC:8623] |
| **ENSG00000018236** | CNTN1 | 1272 | 16 | 2.102580254 | 5.017383551 | 1.405960133 | 3.568652791 | 0.000358822 | contactin 1 [Source:HGNC Symbol;Acc:HGNC:2171] |
| **ENSG00000164853** | UNCX | 340260 | 15 | 19.51546183 | 1.756483252 | 0.368007155 | 4.772959513 | 1.82E-06 | UNC homeobox [Source:HGNC Symbol;Acc:HGNC:33194] |
| **ENSG00000164841** | TMEM74 | 157753 | 14 | 4.85552181 | 3.013458384 | 0.822137796 | 3.665393317 | 0.000246959 | transmembrane protein 74 [Source:HGNC Symbol;Acc:HGNC:26409] |
| **ENSG00000147642** | SYBU | 55638 | 13 | 103.8509184 | 1.419825494 | 0.194878197 | 7.285707259 | 3.2E-13 | syntabulin [Source:HGNC Symbol;Acc:HGNC:26011] |
| **ENSG00000275342** | PRAG1 | 157285 | 12 | 300.7896079 | 0.891127863 | 0.142420997 | 6.256997779 | 3.92E-10 | PEAK1 related, kinase-activating pseudokinase 1 [Source:HGNC Symbol;Acc:HGNC:25438] |
| **ENSG00000183495** | EP400 | 57634 | 11 | 635.4565309 | 0.845401208 | 0.168795553 | 5.008432951 | 5.49E-07 | E1A binding protein p400 [Source:HGNC Symbol;Acc:HGNC:11958] |
| **ENSG00000128713** | HOXD11 | 3237 | 10 | 6.161741011 | 1.917847213 | 0.805140899 | 2.382001976 | 0.017218805 | homeobox D11 [Source:HGNC Symbol;Acc:HGNC:5134] |
| **ENSG00000056291** | NPFFR2 | 10886 | 9 | 29.09294225 | 0.899811384 | 0.244428222 | 3.681290877 | 0.000232056 | neuropeptide FF receptor 2 [Source:HGNC Symbol;Acc:HGNC:4525] |
| **ENSG00000082175** | PGR | 5241 | 8 | 37.11208819 | 1.193295139 | 0.311344267 | 3.832719164 | 0.000126735 | progesterone receptor [Source:HGNC Symbol;Acc:HGNC:8910] |
| **ENSG00000004487** | KDM1A | 23028 | 7 | 9.711064788 | 2.351736019 | 0.603116603 | 3.899305717 | 9.65E-05 | lysine demethylase 1A [Source:HGNC Symbol;Acc:HGNC:29079] |
| **ENSG00000095752** | IL11 | 3589 | 6 | 4417.107715 | 0.845635048 | 0.199191197 | 4.245343485 | 2.18E-05 | interleukin 11 [Source:HGNC Symbol;Acc:HGNC:5966] |
| **ENSG00000167302** | TEPSIN | 146705 | 5 | 160.0140229 | 0.72492128 | 0.198945938 | 3.643810414 | 0.000268631 | TEPSIN adaptor related protein complex 4 accessory protein [Source:HGNC Symbol;Acc:HGNC:26458] |
| **ENSG00000129991** | TNNI3 | 7137 | 4 | 10.14967577 | 1.66230987 | 0.84045026 | 1.977880129 | 0.047942231 | troponin I3, cardiac type [Source:HGNC Symbol;Acc:HGNC:11947] |
| **ENSG00000205795** | CYS1 | 192668 | 3 | 243.1573982 | 0.666973049 | 0.188509173 | 3.538146385 | 0.000402947 | cystin 1 [Source:HGNC Symbol;Acc:HGNC:18525] |
| **ENSG00000178982** | EIF3K | 27335 | 2 | 7248.019048 | 0.707772976 | 0.165627218 | 4.273289041 | 1.93E-05 | eukaryotic translation initiation factor 3 subunit K [Source:HGNC Symbol;Acc:HGNC:24656] |
| **ENSG00000133134** | BEX2 | 84707 | 1 | 122.2473932 | 0.616108683 | 0.159963273 | 3.85156338 | 0.000117366 | brain expressed X-linked 2 [Source:HGNC Symbol;Acc:HGNC:30933] |
